# Top-down modulation in canonical cortical circuits with short-term plasticity

**DOI:** 10.1101/2023.06.13.544791

**Authors:** Felix Waitzmann, Yue Kris Wu, Julijana Gjorgjieva

**Affiliations:** School of Life Sciences, Technical University of Munich, Freising, Germany; Max Planck Institute for Brain Research, Frankfurt, Germany

## Abstract

Cortical dynamics and computations are strongly influenced by diverse GABAergic interneurons, including those expressing parvalbumin (PV), somatostatin (SST), and vasoactive intestinal peptide (VIP). Together with excitatory (E) neurons, they form a canonical microcircuit and exhibit counterintuitive nonlinear phenomena. One instance of such phenomena is response reversal, whereby SST neurons show opposite responses to top-down modulation via VIP depending on the presence of bottom-up sensory input, indicating that the network may function in different regimes under different stimulation conditions. Combining analytical and computational approaches, we demonstrate that model networks with multiple interneuron subtypes and experimentally identified short-term plasticity mechanisms can implement response reversal. Surprisingly, despite not directly affecting SST and VIP activity, PV-to-E short-term depression has a decisive impact on SST response reversal. We show how response reversal relates to inhibition stabilization and the paradoxical effect in the presence of several short-term plasticity mechanisms demonstrating that response reversal coincides with a change in the indispensability of SST for network stabilization. In summary, our work suggests a role of short-term plasticity mechanisms in generating nonlinear phenomena in networks with multiple interneuron subtypes and makes several experimentally testable predictions.

## Introduction

Inhibitory neurons in the cortex are highly diverse in anatomy, electrophysiology, and function (Pfeffer et al., 2013; Kepecs and Fishell, 2014; Jiang et al., 2015; Tremblay et al., 2016). In the mouse cortex, three major classes of interneurons expressing parvalbumin (PV), somatostatin (SST), and vasoactive intestinal peptide (VIP) make up more than 80% of GABAergic interneurons (Tremblay et al., 2016). Together with excitatory (E) neurons, they form a canonical microcircuit relevant for various cortical computations, including locomotion-induced gain modulation (Fu et al., 2014), selective attention (Zhang et al., 2014), context-dependent modulation (Kuchibhotla et al., 2017; Keller et al., 2020), predictive processing (Keller et al., 2012; Attinger et al., 2017), novelty detection (Garrett et al., 2020, 2023), flexible routing of information flow (Yang et al., 2016; Wang and Yang, 2018), regulating global coherence (Veit et al., 2017, 2022), and gating of synaptic plasticity (Canto-Bustos et al., 2022). Interactions between different cell types in the canonical microcircuit can give rise to counterintuitive nonlinear phenomena. More specifically, in darkness, locomotion-induced top-down modulation via VIP decreases the activity of SST neurons in layer 2/3 of mouse primary visual cortex (Fu et al., 2014; Fig. 1). In contrast, when animals receive visual stimuli, locomotion leads to an increase in SST activity (Pakan et al., 2016; Dipoppa et al., 2018; Fig. 1). This phenomenon in which the same locomotion-induced top-down modulation via VIP affects SST response oppositely depending on the visual stimulation condition is known as *response reversal* (Garcia del Molino et al., 2017).

**Fig. 1.**
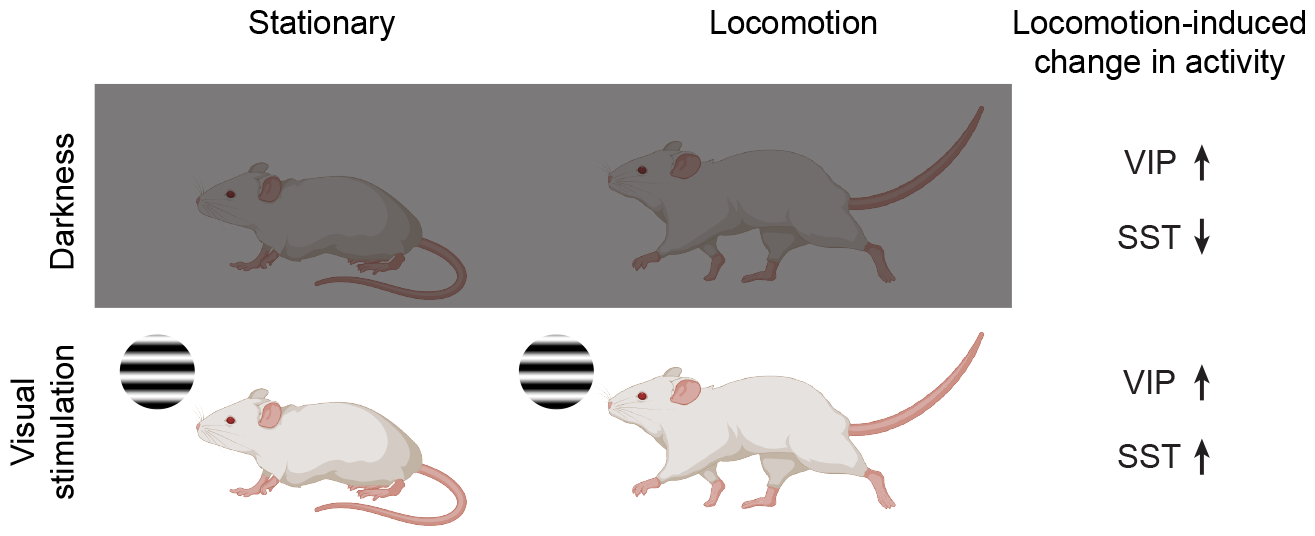
Schematic diagrams illustrating that under different stimulus conditions, locomotion-induced modulatory input via VIP affects SST response oppositely. Top: in darkness, locomotion-induced top-down modulation increases VIP activity but decreases SST activity (Fu et al., 2014). Bottom: in the presence of visual stimulation, locomotion-induced top-down modulation increases both VIP and SST activity (Pakan et al., 2016; Dipoppa et al., 2018).

Previous computational work has shown that networks with nonlinear neuronal input-output functions can generate response reversal (Garcia del Molino et al., 2017). However, cortical neurons exhibit highly irregular spiking (Shadlen and Newsome, 1998) and heterogeneous firing rates (Roxin et al., 2011; Buzsáki and Mizuseki, 2014) that are hallmarks of tightly balanced networks in which population-averaged responses are linear in the input (van Vreeswijk and Sompolinsky, 1998). This raises the possibility that other factors, such as dynamically changing synapses, may contribute to nonlinear population responses like response reversal. On a perceptually and behaviorally relevant timescale from milliseconds to seconds, synapses are subject to short-term plasticity (STP) (Zucker and Regehr, 2002; Markram et al., 2015). Different types of synapses can experience different degrees of short-term depression (STD) or short-term facilitation (STF) (Zucker and Regehr, 2002). In particular, inhibitory synapses exhibit more pronounced short-term dynamics than excitatory synapses, and synapses from different interneuron subtypes can undergo different short-term plastic changes (Campagnola et al., 2022). However, little is known about how these experimentally identified short-term plasticity mechanisms shape network dynamics and computations in recurrent neural circuits of multiple interneuron subtypes.

Response reversal of SST induced by the same top-down modulation may suggest that the network operates in different regimes under different stimulation conditions. Increasing evidence suggests that cortical networks operate in an inhibition-stabilized regime, in which feedback inhibition generated by the network is imperative to stabilize excitatory activity (Tsodyks et al., 1997; Sanzeni et al., 2020). In networks with one excitatory and one inhibitory population and fixed connectivity, an identifying characteristic of inhibition stabilization is that increasing (decreasing) excitatory input to the inhibitory population decreases (increases) inhibitory firing, known as the paradoxical effect (Tsodyks et al., 1997; Li et al., 2019; Miller and Palmigiano, 2020). Yet, it is unclear whether response reversal can be linked to inhibition stabilization and whether there exists a relationship between response reversal and the paradoxical effect. In addition, how short-term plasticity shapes inhibition stabilization in networks with multiple interneuron subtypes, particularly how specific interneuron subtypes contribute to network stabilization (which we refer to as *interneuron-specific stabilization*), is unknown.

Here, we use analytical calculations and numerical simulations to demonstrate that inhibitory short-term plasticity enables response reversal without requiring neuronal nonlinearities. We find that despite not directly affecting SST and VIP activity, PV-to-E STD has a crucial influence on response reversal. We further reveal the relationship between response reversal, the paradoxical effect, and the interneuron-specific stabilization property of the network. Interestingly, when the SST response to top-down modulation switches from suppression to enhancement, the network undergoes an interneuron-specific change in stabilization, and SST is required for network stabilization. In summary, our model suggests that inhibitory short-term plasticity enables the network to perform nonlinear computations and makes several experimentally testable predictions.

## Results

To study how response reversal emerges in canonical cortical circuits, we used rate-based population models consisting of one excitatory (E) and three different inhibitory (PV, SST, VIP) populations with network connectivity constrained by previous experimental studies (Fig. 2A; Pfeffer et al., 2013). This type of model allows for a trade-off between sufficient biological detail and mathematical analysis and has previously been used with great success to study cortical computations (Murphy and Miller, 2009; Litwin-Kumar et al., 2016; Garcia del Molino et al., 2017; Mahrach et al., 2020; Richter and Gjorgjieva, 2022). Consistent with experimental work (Sanzeni et al., 2020), network connectivity was chosen so that the network operates in an inhibition-stabilized regime defined as the regime where feedback inhibition generated by the network is needed to stabilize recurrent excitation (Tsodyks et al., 1997). As proposed by influential modeling work on cortical dynamics (van Vreeswijk and Sompolinsky, 1996, 1998), the network’s population-averaged responses can be approximated by a rectified linear function of the input (Fig. 2A, inset). To account for activity-dependent changes in network connectivity on a perceptually and behaviorally relevant timescale, we modeled short-term plasticity based on recent experimental work from the Allen Institute (Campagnola et al., 2022). We incorporated the four most pronounced short-term plasticity mechanisms: PV-to-E short-term depression (STD), PV-to-PV STD, PV-to-VIP STD, and SST-to-VIP short-term facilitation (STF) (Fig. 2A; Fig. S1). Since all the prominent synapses undergoing short-term plasticity are inhibitory, we refer to the plasticity mechanisms as *inhibitory short-term plasticity* (iSTP). The dynamics of the network with iSTP can be described as follows:

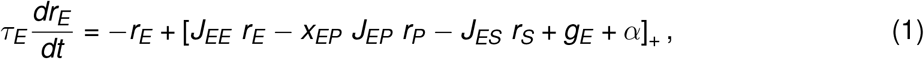

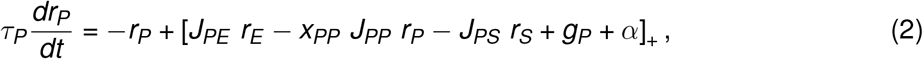

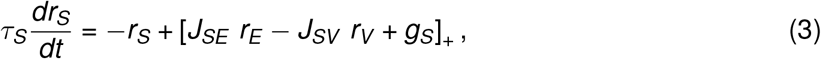

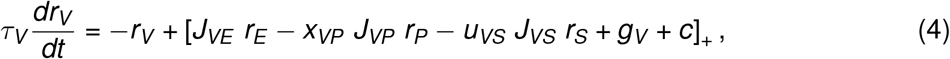

for the rates *r*_*i*_ of excitatory, PV, SST, and VIP populations with *i* ∈ {*E, P, S, V* } and [·]_+_ denotes linear rectification. *τ*_*i*_ represents the corresponding time constant of the rate dynamics, *J*_*ij*_ denotes the synaptic strength from population *j* to population *i*, and *g*_*i*_ is the individual background input. Importantly, we distinguish between bottom-up input to E and PV to represent different stimulation conditions, denoted as *α*, and top-down input to VIP mimicking locomotion-induced top-down modulation, denoted as *c* (Fig. 2A).

**Fig. 2.**
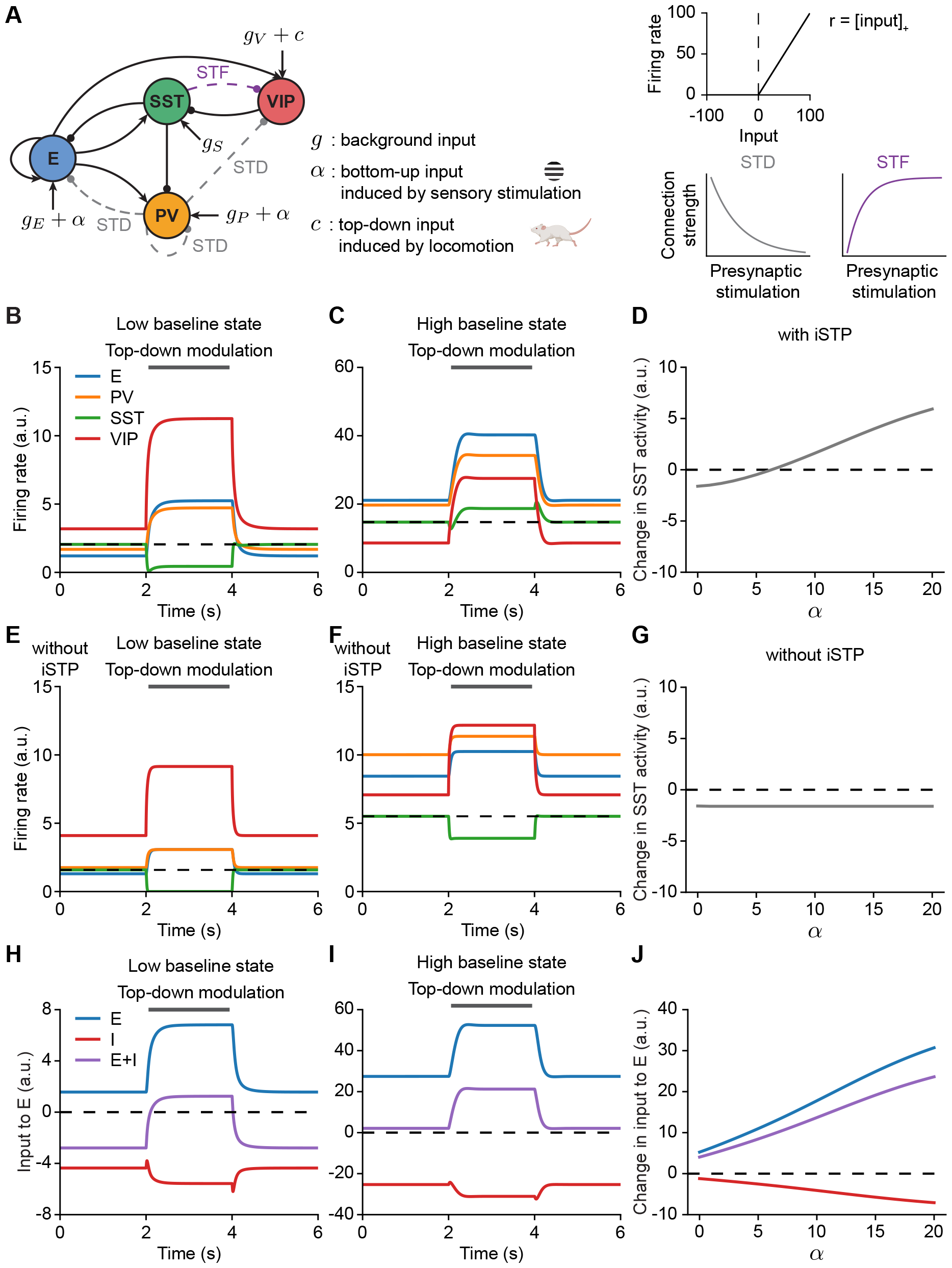
Inhibitory short-term plasticity enables response reversal of SST induced by top-down modulation via VIP. (**A**) Schematic of network model with one excitatory (E) population and three distinct inhibitory populations, including PV, SST, and VIP. Short-term depressing (STD) and short-term facilitating (STF) connections are indicated by the dashed lines. Each population receives a background input *g*. E and PV receive bottom-up input *α* depending on sensory stimulation, and VIP receives top-down input *c* during locomotion. Top right: rectified linear input-output function; Bottom right: cartoons showing how inhibitory connection strength changes with presynaptic stimulation under STD and STF. (**B**) Network responses to top-down modulation without any bottom-up input (*α* = 0), corresponding to darkness without sensory stimulation. Top-down modulation via VIP is applied during the interval from 2 to 4 s (gray bar). Different colors denote the activity of different populations. The dashed line represents the initial activity level of SST. (**C**) Same as B but at *α* = 15 corresponding to sensory stimulation. (**D**) Change in SST response induced by top-down modulation to VIP as a function of bottom-up input *α* in networks with iSTP. (**E**) Same as B but for networks without iSTP. (**F**) Same as C but for networks without iSTP. (**G**) Same as D but for networks without iSTP. (**H**) Input to the E population at *α* = 0. Different colors indicate different sources: input from the E population, input from the I populations, and the sum of the inputs from the E and I populations. (**I**) Same as E but at *α* = 15. (**J**) Change in different sources of recurrent inputs to the E population measured between baseline and at steady state during top-down modulation as a function of bottom-up input *α*.

Short-term plasticity mechanisms are implemented based on the Tsodyks-Markram model (Tsodyks et al., 1998):

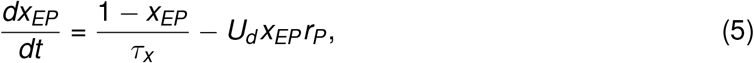

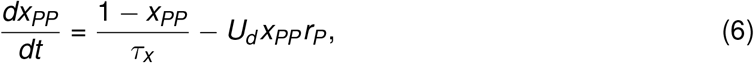

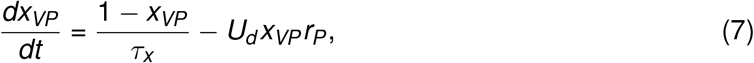

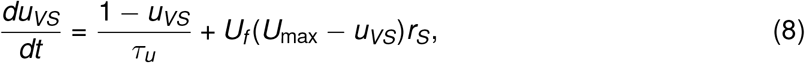

where *x*_*ij*_ is a short-term depression variable limited to the interval (0,1] for the synaptic connection from population *j* to population *i*. Biophysically, the short-term depression variable *x* represents the fraction of vesicles available for release. *τ*_*x*_ is the time constant of STD, and *U*_*d*_ is the depression factor controlling the degree of depression induced by the presynaptic activity. Similarly, *u*_*ij*_ is a short-term facilitation variable constrained to the interval [1, *U*_*max*_) for the synaptic connection from population *j* to population *i*. Unlike short-term depression, the short-term facilitation variable *u* biophysically represents the ability to release neurotransmitters. *τ*_*u*_ is the time constant of STF, *U*_*f*_ is the facilitation factor controlling the degree of facilitation induced by the presynaptic activity, and *U*_*max*_ is the maximal facilitation value.

### iSTP enables response reversal of SST

To represent different stimulus conditions (e.g., darkness vs. visual stimulation), we varied the bottom-up input *α* to E and PV. Increasing *α* leads to a supralinear increase in the baseline activity in all populations (Fig. S2). We modeled the effect of locomotion-induced top-down modulation on network activity by increasing the input to VIP by a positive value *c*. In our network model with iSTP, for a low *α*, corresponding to low baseline activity in the absence of bottom-up input, top-down modulation via additional excitatory input to VIP decreases SST activity (Fig. 2B). In contrast, for a high *α* corresponding to high baseline activity, the same top-down modulation leads to an increase in SST activity and, thus, response reversal (Fig. 2C). Our modeling results suggest that under different stimulus conditions regulated by bottom-up inputs, identical top-down modulation reversely affects the change of SST activity (Fig. 2D), consistent with previous experiments (Fu et al., 2014; Pakan et al., 2016; Dipoppa et al., 2018).

To highlight the role of iSTP in generating response reversal of SST activity, we further simulated the same network while disabling iSTP (Fig. 2E, F). In contrast to networks with iSTP, the change of SST activity is largely unaffected for different values of *α* when iSTP is disabled (Fig. 2G). Interestingly, in our model, for a high *α* during the stimulation period, despite the increased activity of all inhibitory populations, the steady state of excitatory activity also increases (Fig. 2C). This observation appears to differ from what would be predicted by a classical disinhibition mechanism in which reducing inhibition increases excitatory activity. We confirm this by plotting the amount of recurrent excitation, recurrent inhibition, and the sum of recurrent excitation and inhibition that the excitatory population receives during the simulation(Fig. 2H, I). Surprisingly, even for a low *α*, despite decreased SST activity, top-down modulation via VIP increases the total inhibition to the excitatory population at the steady state (Fig. 2H). Enhanced inhibition to the excitatory population at the steady state during top-down modulation is also observed for a high *α* (Fig. 2I). We systematically investigated the change in the input to the excitatory population due to top-down modulation at different levels of bottom-up input *α*. We found that top-down modulation always increases the amount of inhibition to the excitatory population irrespective of whether it increases or decreases the activity of the SST population as *α* changes (Fig. 2J). These results suggest that rather than the decrease in the total inhibition, the increase in the recurrent excitation contributes to the elevated excitatory activity (Fig. 2H-J). Importantly, a joint increase in the excitation and inhibition of the excitatory population is a distinctive feature in inhibition-stabilized networks (Litwin-Kumar et al., 2016; Miller and Palmigiano, 2020; Wu and Gjorgjieva, 2023). In contrast, non-inhibition-stabilized networks do not exhibit an increase in the total inhibitory inputs to the excitatory population (Fig. S3).

Taken together, our numerical simulation results reveal that experimentally identified forms of iSTP enable response reversal, a nonlinear computation observed in cortical circuits.

### Theoretical analysis

To better understand how iSTP enables our model network to perform response reversal of SST activity, we sought to mathematically analyze how top-down modulation affects SST activity. Locomotion-induced top-down modulation can be considered a form of perturbation to the VIP population. Investigating how top-down modulation inversely affects SST activity under different stimulus conditions can therefore be mathematically formulated as how perturbations to VIP affect SST activity under varying levels of *α*. To this end, we extended previous studies on static networks (Garcia del Molino et al., 2017; Palmigiano et al., 2023) and developed a general theoretical framework for networks with iSTP. More specifically, we derived how the steady state response of any population changes with a perturbation of the external input to a given population while including iSTP (see Methods). Using this approach, we formulated the change of SST activity induced by top-down modulation via the change of the input to VIP, **R**_*SV*_, as follows:

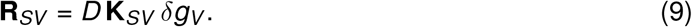

Here, *D* is a positive quantity for any stable network (see Methods, Fig. S4A), *δg*_*V*_ represents the perturbation of the VIP population’s input which is a positive number *c* in our network setting, and **K**_*SV*_ is the response factor which is given by:

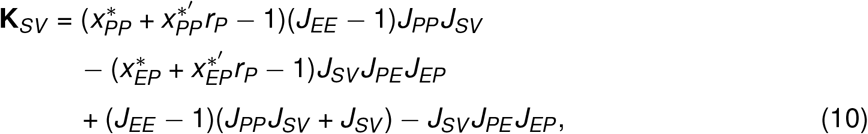

with 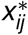 the short-term plasticity variable from population *j* to population *i* at steady state before perturbation, and 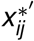 the derivative of the short-term plasticity variable with respect to the activity of population *j*, evaluated at the steady state (see SI Text).

A negative (positive) **R**_*SV*_ denotes a decrease (increase) in SST activity caused by top-down modulation. To have response reversal, **R**_*SV*_ must switch its sign for different values of *α*, corresponding to different stimulus conditions. More specifically, when animals perceive no visual stimulus in darkness, i.e., when *α* is low, top-down modulation via VIP decreases SST activity. Therefore, **R**_*SV*_ is expected to be negative for a low *α*. In contrast, when animals receive a visual stimulus, namely when *α* is high, top-down modulation via VIP increases SST activity. Thus, **R**_*SV*_ is expected to be positive for a high *α*.

In agreement with our simulation results (Fig. 2B, C), we observed that **R**_*SV*_ changes its sign when calculated at different values of *α* (Fig. 3). As our theoretical framework is based on the linearization of the network dynamics around the steady state and higher order terms are ignored (see Methods), the computed **R**_*SV*_ agrees well with the numerical simulation results for small perturbations (Fig. 3A) and diverges for large perturbations (Fig. 3B). Yet, it qualitatively captures the key aspect of modeling behaviors: the sign switch of the change in SST activity induced by top-down modulation with different values of *α*. Note that while here we are interested in how neural activity changes in response to a given perturbation, several other studies have investigated the contributions of higher-order motifs to the perturbation-induced change of neural activity in excitatory and inhibitory networks (Pernice et al., 2011; Sadeh and Clopath, 2020).

**Fig. 3.**
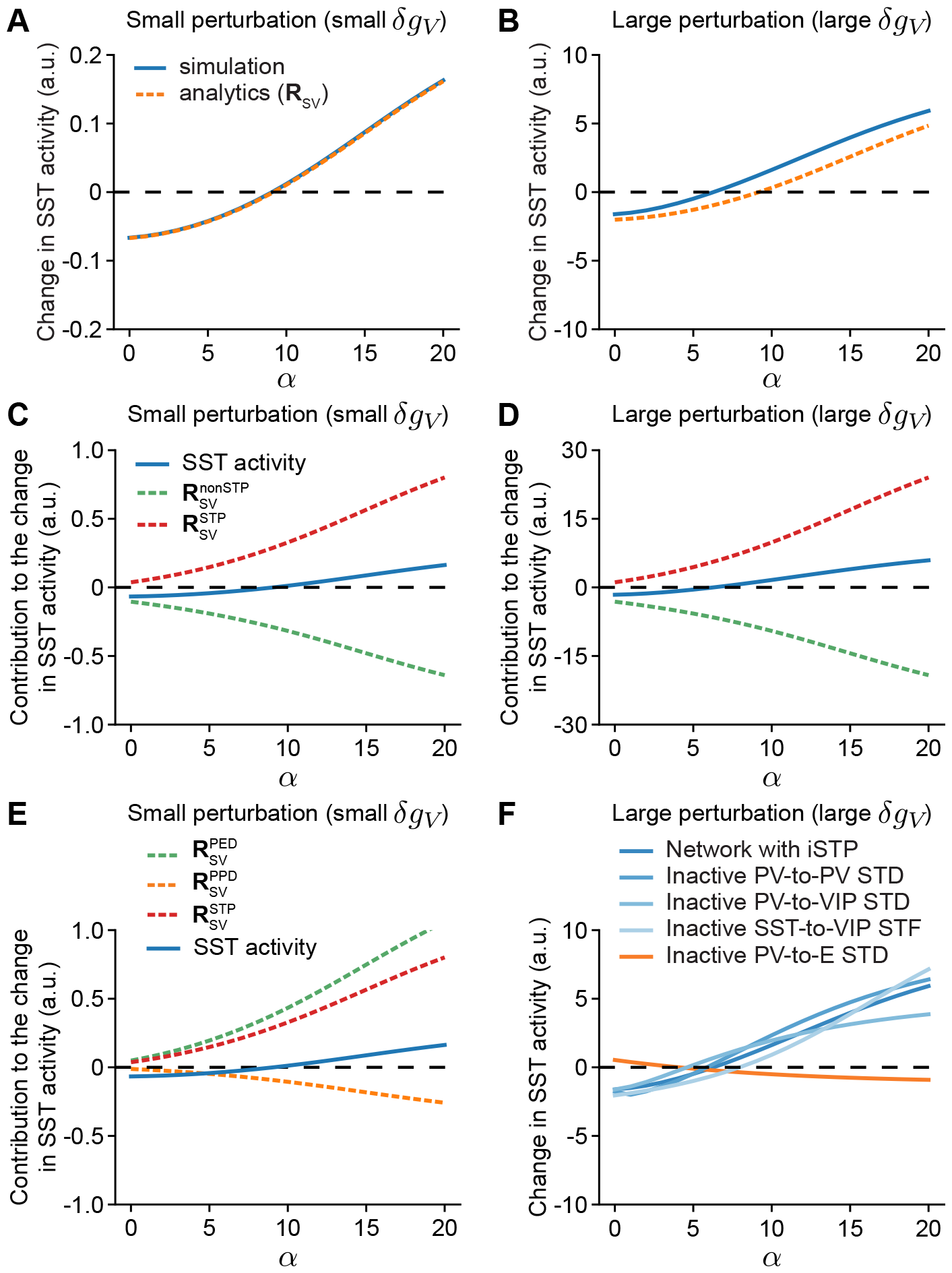
Comparison between analytical predictions and numerical simulations on the change in SST activity and identification of PV-to-E STD as the crucial STP mechanism for the generation of response reversal. (**A**) Analytical prediction of the change in SST population response induced by the perturbation to VIP (**R**_*SV*_) matches closely with numerical simulation for a small perturbation. (**B**) Same as A but with a large perturbation. (**C**) Analytical contributions of the STP-dependent term 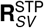 and the STP-independent term 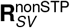 to the change in SST activity as a function of bottom-up input *α* for a small perturbation. (**D**) Same as C but with a large perturbation. (**E**) Analytical contributions of the PV-to-E STD-dependent term 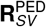, the PV-to-PV STD-dependent term 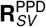, and the overall STD-dependent term 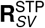to the change in SST activity as a function of bottom-up input *α* for a small perturbation. (**F**) Change in SST response induced by top-down modulation to VIP as a function of bottom-up input *α* with a large perturbation for different network configurations marked with different colors. Here, for small perturbations *δg*_*V*_ = 0.1 and for large perturbations *δg*_*V*_ = 3.

As shown in Eq. 9, since *D* and *δg*_*V*_ are positive, **K**_*SV*_ is the only term that can change the sign of **R**_*SV*_. To further investigate the influence of the iSTP mechanisms on response reversal, we rewrote **K**_*SV*_ as a sum of a short-term plasticity-dependent and a short-term plasticity-independent term:

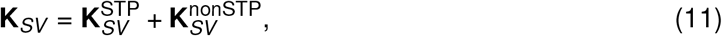

where

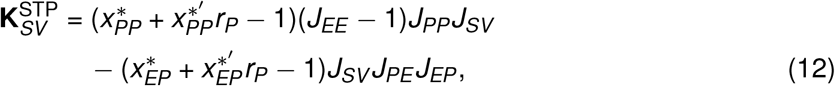

and

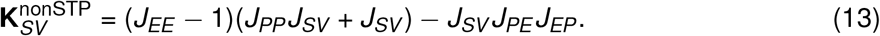

Analogously to **K**_*SV*_, we have:

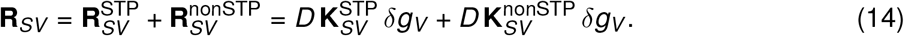

As the short-term plasticity-independent term 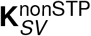is governed by the static network weights, it is constant over the entire range of change in bottom-up input, i.e., 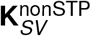does not change with *α*. Note that because of Eq. 9 and since *D* is always positive but subject to change in magnitude (Fig. S4A), 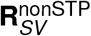 changes in magnitude as well (Fig. 3B). To match recent experimental findings that the network is inhibition stabilized when animals receive no stimulus in darkness (Sanzeni et al., 2020), we set *J*_*EE*_ to be larger than 1. In this case, the short-term plasticity-independent term 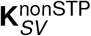 is always negative, which implies that 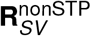 is also always negative. Thus, the short-term plasticity-dependent term 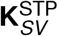, and as a result, 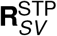 too, is the only part that can influence the sign of **R**_*SV*_ and enable the network to perform response reversal of SST activity (Fig. 3C, D).

In conclusion, our theoretical framework enables us to analyze how perturbations affect the activity of individual populations and hence reveals how iSTP enables response reversal of SST activity.

### PV-to-E STD plays a key role in the generation of response reversal

Next, we sought to dissect the role of individual iSTP mechanisms in response reversal. To this end, we separated the short-term plasticity-dependent term 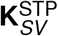 (Eq. 12) into a PV-to-PV STD-dependent part 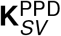 and a PV-to-E STD-dependent part 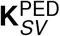as follows:

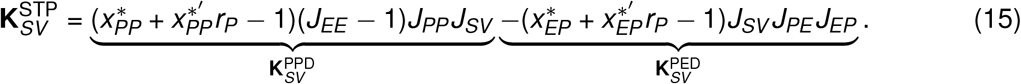

Since both 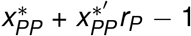 and 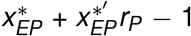 are always negative (see Methods, Fig. S4B), the PV-to-PV STD-dependent part 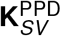 is always negative and the PV-to-E STD-dependent part 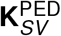 is always positive. Importantly, 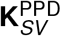 decreases with increasing bottom-up input *α*, whereas 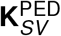 increases with increasing bottom-up input *α* (see SI Text, Fig. S4C).

Similarly, we can write:

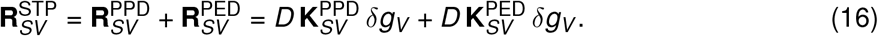

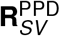 and 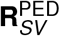 show similar changes as 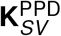 and 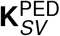, respectively (Fig. 3E). Therefore, when bottom-up input *α* increases from a low to a high level (e.g., switching from darkness to visual stimulation condition), to display response reversal 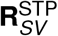 must overcome in magnitude the negative 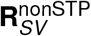, resulting in an overall switch of **R**_*SV*_ from negative to positive. The increasing PV-to-E STD-dependent term rather than the decreasing PV-to-PV STD-dependent term is imperative for this switch (Fig. 3E). As no terms directly associated with PV-to-VIP STD and SST-to-VIP STF appear in Eq. 15, our analysis shows that these two mechanisms are unimportant for the generation of response reversal.

We performed the same simulations as with the intact network while inactivating specific iSTP mechanisms to confirm our analysis. The inactivation of particular mechanisms was implemented by freezing the respective plasticity variables at their baseline values when bottom-up input is high, ensuring that the steady-state activities of all populations are positive at the baseline and during the top-down modulation period. We then varied the bottom-up inputs from high to low and found that SST response reversal still occurs despite inactivating PV-to-PV STD, PV-to-VIP STD, or SST-to-VIP STF. Such networks show similar patterns to networks with intact iSTP (Fig. 3F). In contrast, when PV-to-E STD is inactivated, the change of SST activity manifests an opposite trend from that in networks with intact iSTP (Fig. 3F). Furthermore, we found that PV-to-E STD is crucial for generating the effective supralinear input-output relation observed in the baseline state for varying bottom-up input *α* (Fig. S2). Inactivating PV-to-E STD completely diminished the supralinearity of the effective input-output relations in contrast to inactivating other iSTP mechanisms (Fig. S5A). In addition, as bottom-up input increases, the resulting inhibitory current from PV to E is suppressed by PV-to-E STD (Fig. S5B). This suppression is greater for stronger bottom-up inputs leading to a sublinear increase in PV current, which is important for the generation of the effective supralinear input-output relation (Fig. S5B).

Taken together, our analysis and numerical simulations reveal that PV-to-E STD is the determining mechanism for generating response reversal. In contrast, the effects of PV-to-PV STD, PV-to-VIP STD, and SST-to-VIP STF on response reversal are negligible.

### Relationship between response reversal and the paradoxical effect

Locomotion-induced top-down modulation excites VIP and effectively inhibits SST due to the mutually inhibitory connections between VIP and SST. However, when animals receive visual stimuli at a high baseline activity state (high *α*), additional VIP inhibition induced by top-down modulation increases the activity of SST. This phenomenon is reminiscent of the paradoxical effect (Tsodyks et al., 1997; Ozeki et al., 2009). We thus sought to identify the relationship between response reversal and the paradoxical effect. To this end, we derived the change of SST activity induced by a change in the input to SST itself, **R**_*SS*_, as follows:

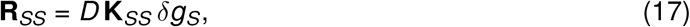

where

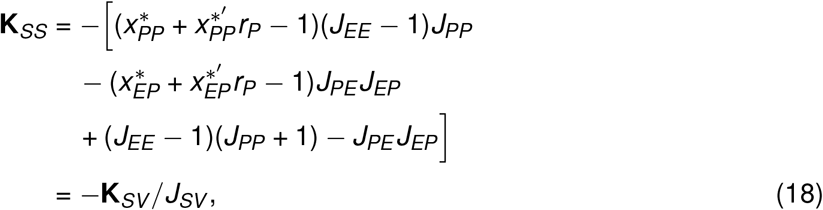

and *δg*_*S*_ represents the change of input to SST. Furthermore,

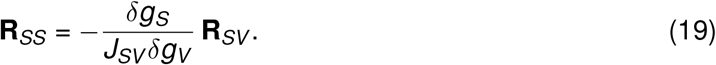

When *δg*_*S*_ is positive, to obtain a paradoxical response of SST (i.e., to have a negative **R**_*SS*_), **K**_*SS*_ has to be negative. As **K**_*SS*_ is equal to −**K**_*SV*_ */J*_*SV*_, for low *α* corresponding to the darkness condition (**K**_*SV*_ and **R**_*SV*_ are negative), **K**_*SS*_ and **R**_*SS*_ are positive, hence, no paradoxical response is observed (Fig. 4A, B left). In contrast, for high *α* corresponding to the visual stimulation condition (**K**_*SV*_ and **R**_*SV*_ are positive), **K**_*SS*_ and **R**_*SS*_ are negative. Therefore, SST exhibits a paradoxical response (Fig. 4A, B right).

**Fig. 4.**
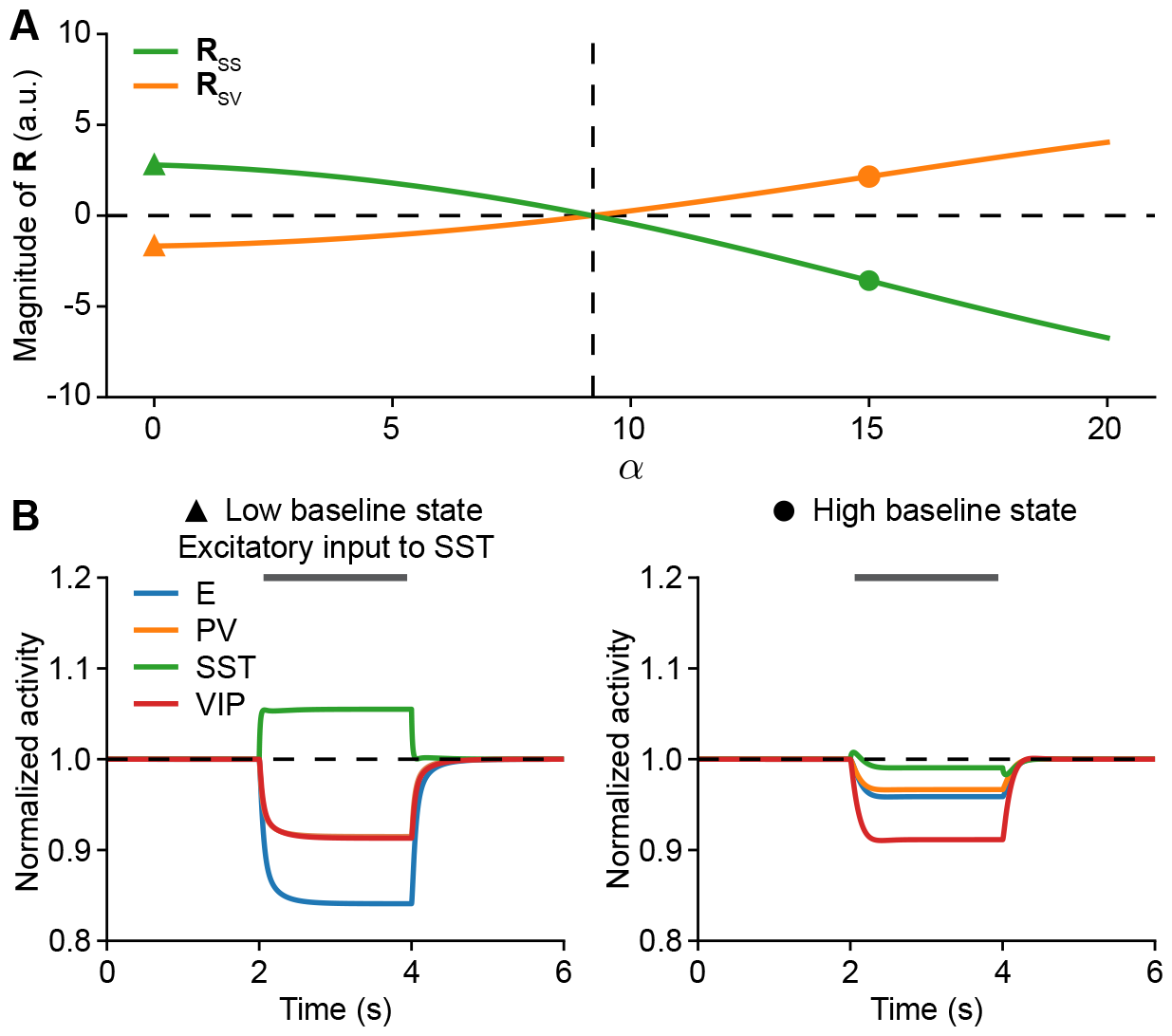
Relationship between response reversal and paradoxical response of SST. (**A**) Analytical predictions of the change in SST response induced by an excitatory perturbation (*δg*_*S*_) to SST, **R**_*SS*_, and change in SST response induced by an excitatory perturbation (*δg*_*V*_) to VIP, **R**_*SV*_, as a function of bottom-up input *α*. Here, *δg*_*V*_ = *δg*_*S*_ = 3. (**B**) Left: Normalized activity when injecting additional excitatory current into SST at a low baseline state corresponding to *α* = 0 marked with triangular in A. SST does not show a paradoxical response. Right: Same as left but at a high baseline state corresponding to *α* = 15 marked with a dot in A. SST shows a paradoxical response.

We have mathematically proven a correspondence between response reversal and the paradoxical response of SST. More specifically, the SST population will not show a paradoxical response when top-down modulation via VIP decreases SST activity, but will respond paradoxically when top-down modulation via VIP increases SST activity.

### Relationship between response reversal, the paradoxical effect, and interneuron-specific stabilization

The paradoxical effect is a defining characteristic of inhibition stabilization in networks with fixed connectivity (Tsodyks et al., 1997). We, therefore, sought to investigate the relationship between response reversal and inhibition stabilization. Identifying the relationship may shed light on how response reversal relates to other cortical functions as inhibition-stabilized networks can perform a variety of computations (Sadeh and Clopath, 2021). As the network in our study consists of three different inhibitory populations, the network can, in principle, be stabilized by any type of interneuron. Beyond identifying inhibition stabilization, we particularly aimed to ascertain the specific interneuron subtype that stabilizes the model networks in different stimulation conditions.

To this end, we computed the leading eigenvalue of the Jacobian of individual subnetworks with the corresponding firing rates and short-term plasticity dynamics while excluding specific interneuron subtypes. Such eigenvalues can be used to determine the stability of the subnetwork. A negative leading eigenvalue implies that the fixed point of the network dynamics is stable and a transient perturbation to the system does not result in a deviation from the original fixed point. In contrast, a positive leading eigenvalue means that the fixed point is unstable, and a transient perturbation causes a deviation from the original fixed point. We found that the leading eigenvalue of the Jacobian of the E subnetwork in the model (defined as the network without any interneurons) is positive for all values of *α*, suggesting that the E subnetwork is unstable and the network is inhibition-stabilized for all stimulation conditions (Fig. S6). Furthermore, we found that the E subnetwork being unstable (i.e. *J*_*EE*_ *>* 1) at the high bottom-up input is a necessary condition to observe response reversal (see SI Text). VIP does not stabilize the network, as the leading eigenvalue of the Jacobian of the E-VIP subnetwork (the network without PV and SST interneurons) is always positive (Fig. S6). By computing the leading eigenvalue of the Jacobian of the E-PV-VIP subnetwork (the network without SST interneurons), we found that the eigenvalue switches from negative to positive when the response of SST to top-down modulation is reversed (Fig. 5A), indicating that SST is required for network stabilization when top-down modulation via VIP increases SST activity. Furthermore, the leading negative eigenvalue of the Jacobian of the E-PV-VIP sub-network for low *α* suggests that in the regime in which top-down modulation via VIP decreases SST activity, the network does not require SST for stabilization and can be stabilized by PV. To determine whether PV could be the only interneuron subtype stabilizing the network in that regime, we calculated the leading eigenvalue of the Jacobian of the E-SST-VIP subnetwork (the network without PV interneurons). We found that this eigenvalue is always negative in the current model (Fig. 5A), suggesting that SST can serve the stabilization role in that regime as well as PV.

**Fig. 5.**
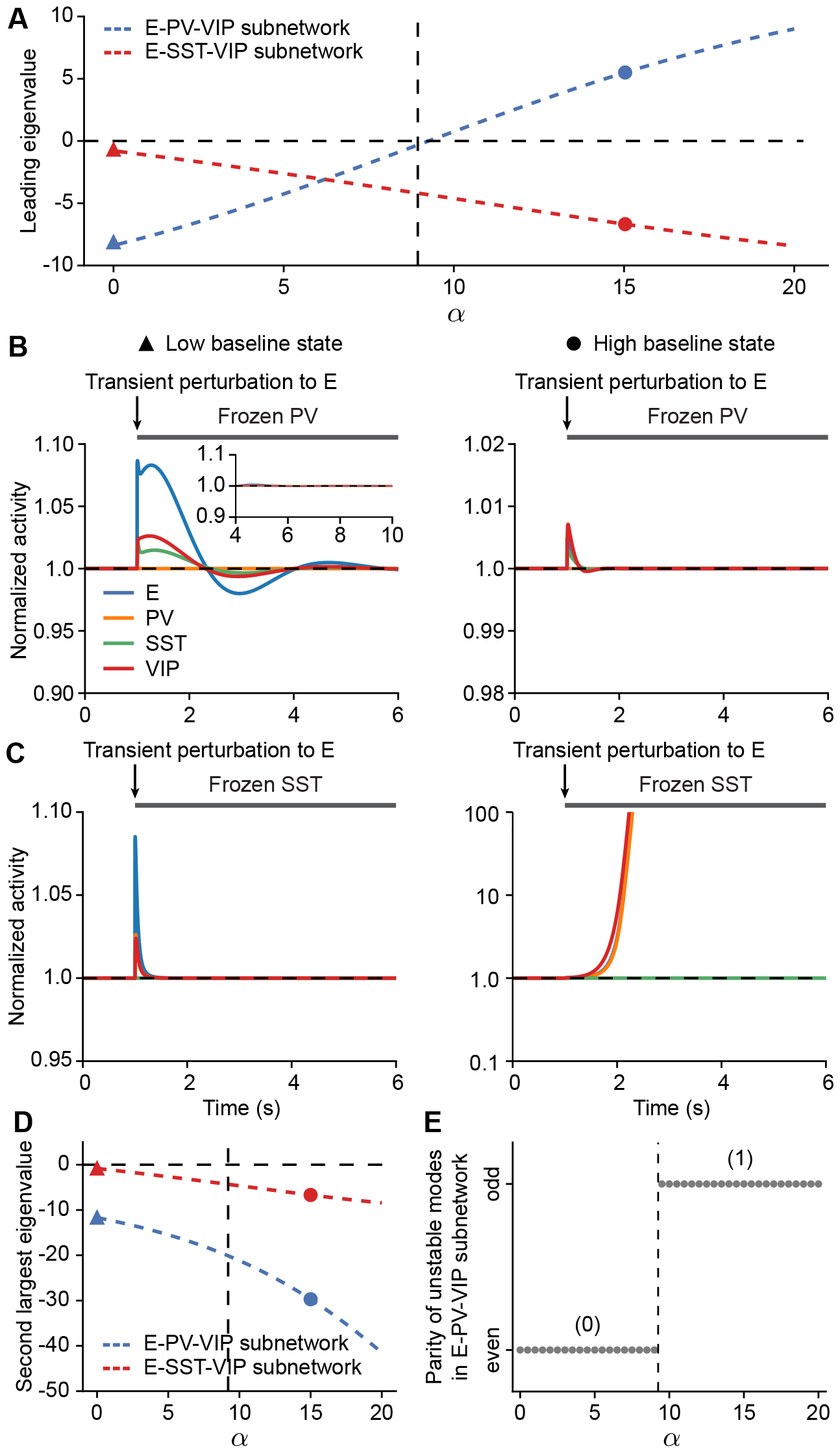
The network undergoes a change in the indispensability of SST for network stabilization with increasing bottom-up input. (**A**) Leading eigenvalues of the E-PV-VIP subnetwork and the E-SST-VIP subnetwork as a function of bottom-up input *α*. The response reversal boundary extracted from analytical calculations (**R**_*SV*_ = 0) is indicated by the vertical dashed line. (**B**) Left: Normalized activity when injecting an additional transient excitatory current into E while freezing PV for networks at a low baseline state corresponding to *α* = 0 marked with a triangle in A. The small transient excitatory input is introduced at the time marked with arrows. The periods in which PV is frozen are marked with the gray bar. Right: Same as left but for networks at a high baseline state corresponding to *α* = 15 marked with a dot in A. (**C**) Similar to B but with frozen SST. (**D**) Same as A but for the second largest eigenvalue. (**E**) Parity of the number of unstable modes in the E-PV-VIP subnetwork as a function of bottom-up input *α*. Numbers indicate the amount of unstable modes.

**Fig. 6.**
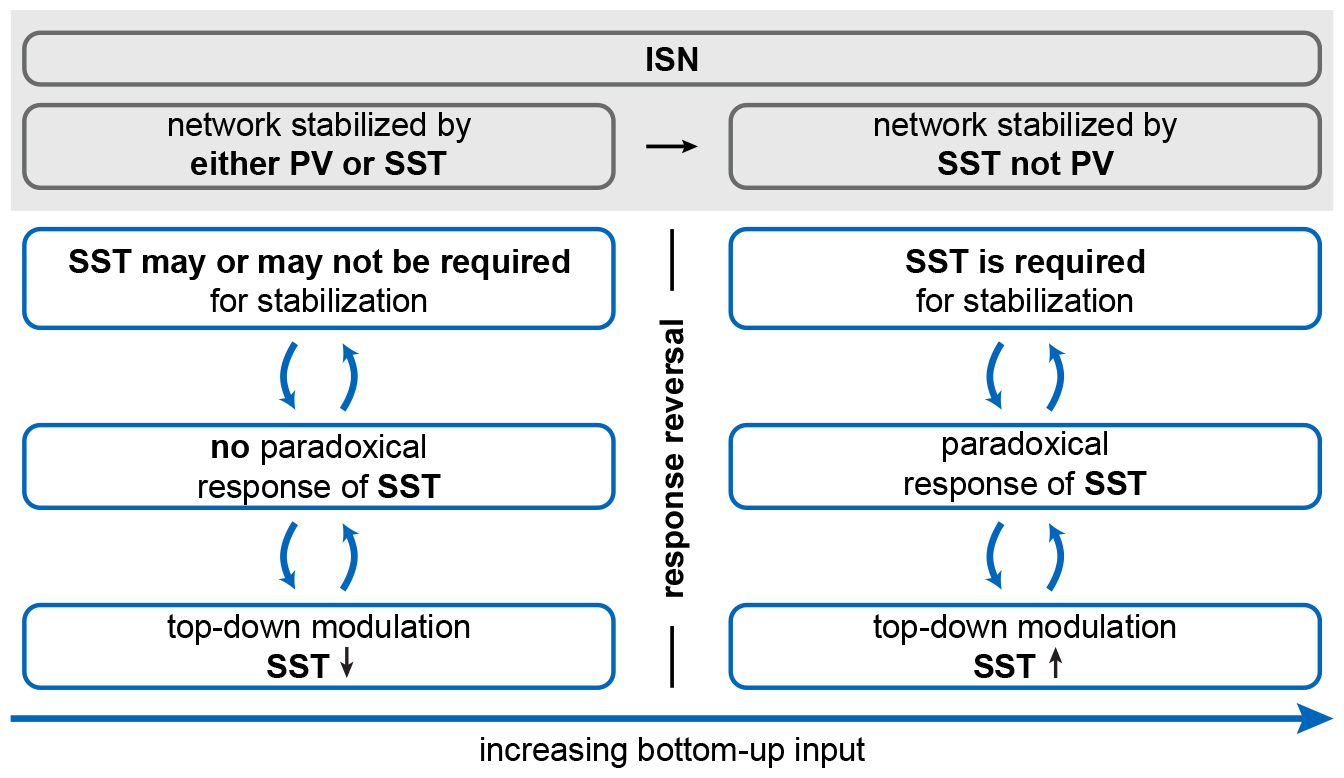
The relationship between response reversal, paradoxical effect, and inhibition stabilization. At low bottom-up input, top-down modulation decreases SST activity, and the network does not exhibit a paradoxical response of SST such that SST may not be required for stabilization, or SST may be required for stabilization, but the E-PV-VIP subnetwork has an even number of unstable modes. As demonstrated in Fig. 5, in this regime, the network is inhibition stabilized and stabilized by either PV or SST. With increasing bottom-up input, the response of SST induced by top-down modulation is reversed from suppression to enhancement, the network exhibits a paradoxical response of SST and requires SST for stabilization with an odd number of unstable modes in the E-PV-VIP subnetwork. As demonstrated in Fig. 5, in this regime, the network is inhibition stabilized and stabilized by SST but not PV. Note that while the relationship between response reversal, paradoxical effects, and inhibition stabilization marked in blue boxes does not depend on the choice of parameters, the possible interneuron-specific stabilization regimes shaded in gray are contingent on specific parameters (see SI Text).

We confirmed these results by injecting a transient excitatory perturbation into the excitatory population while clamping the activity of either PV or SST. We found that when clamping PV activity, the fixed point in the given network is stable to perturbations over the entire range of *α* and reaches the same fixed point after the transient perturbation (Fig. 5B). In contrast, when clamping SST activity, while the fixed point at low *α* is stable to perturbations, a transient perturbation at high *α* leads to unstable dynamics (Fig. 5C).

Consistent with the change in the requirement of SST for network stabilization, we observed a transition in the prevalence of inhibition received by the excitatory population from PV to SST with increasing *α* (Fig. S7). More specifically, at the low baseline state, the excitatory population receives more inhibition from PV than SST (Fig. S7A). Top-down modulation via VIP leads to increases in the overall inhibition and the inhibition from PV at the steady state but a decrease in the inhibition from SST (Fig. S7A). In contrast, at the high baseline state, the excitatory population receives more inhibition from SST than PV (Fig. S7B). Top-down modulation increases the overall inhibition at the steady state as well as the inhibition from both PV and SST (Fig. S7B). This increase in total inhibition at the steady state observed during top-down modulation is a unique characteristic of inhibition-stabilized networks in contrast to non-inhibition-stabilization networks (Fig. S3; Litwin-Kumar et al., 2016; Miller and Palmigiano, 2020; Wu and Gjorgjieva, 2023). In inhibition-stabilized networks with iSTP, top-down modulation induces a transient disinhibition enabling the growth of recurrent excitation and increasing excitation and inhibition to the excitatory population at the steady state.

To systematically investigate how response reversal and paradoxical effects of SST relate to interneuron-specific stabilization, we conducted mathematical analyses and found that **K**_*SV*_ and **K**_*SS*_ are linked to the determinant of the Jacobian of the E-PV-VIP network, det(**M**_E-PV-VIP_) (see SI Text). In the network we considered here, because of the short-term plasticity mechanisms, the Jacobian of the E-PV-VIP subnetwork is a 6-by-6 matrix. When **K**_*SV*_ is positive (i.e., top-down modulation increases SST activity), **K**_*SS*_ is negative (i.e., the network exhibits paradoxical effects of SST), and det(**M**_E-PV-VIP_) is negative. Note that in high-dimensional systems, the determinant of the Jacobian matrix alone is not sufficient to determine network stability. For a six-dimensional system, a negative det(**M**_E-PV-VIP_) implies an odd number of positive eigenvalues corresponding to unstable eigenvectors/modes and thus the necessity for SST stabilization. However, the network can also require SST for stabilization in the presence of a positive det(**M**_E-PV-VIP_), for instance, when the Jacobian of the E-PV-VIP subnetwork has an even number of unstable modes. To confirm the change in the number of unstable modes, we examined the second-largest eigenvalue of the E-PV-VIP subnetwork and found that the second-largest eigenvalue is always negative in the given network (Fig. 5D). As a result, the number of unstable modes changes from even to odd (Fig. 5E) when the SST response reverses from suppression to enhancement, and the network exhibits the paradoxical effect in the response of SST. Note that we did not find a direct mathematical relationship between PV stabilization and response reversal of SST (see SI Text). In other words, response reversal does not imply a change in PV stabilization. Consequently, PV stabilization and how it changes with bottom-up inputs, as presented in our study, are contingent on specific parameters.

Taken together, these results suggest that with increasing bottom-up input, representing a change in stimulation condition, the impact of top-down modulation on SST activity transitions from suppression to enhancement, the network exhibits a paradoxical response of SST, requires SST for stabilization, and the E-PV-VIP subnetwork has an odd number of unstable modes (Figs. 4A, 5A, 6).

### Modeling results are robust to variations in short-term plasticity mechanisms, inputs, and network connectivity

To demonstrate that our results are valid for a variety of perturbations, we performed different sensitivity analyses on short-term plasticity mechanisms, inputs, and network connectivity.

We first investigated if additional short-term plasticity mechanisms affect our results. In this study, we used a rate-based population model, ignoring the large number of connections between individual neurons on a microscopic level. Given the dominant number of excitatory neurons in the cortex, we might have underestimated the effective depression of the E-to-E connection and facilitation of the E-to-SST connection compared to real circuits (Campagnola et al., 2022). We therefore sought to examine their influence on our results by analyzing how they might affect the analytical expression of **K**_*SV*_ and network simulations. We found that the response reversal of SST from suppression to enhancement with increasing bottom-up input, as reported experimentally, is preserved in the presence of E-to-E STD (Fig. 7A). However, the change in SST activity evolves non-monotonically with increasing bottom-up input, starting to decrease and eventually being reversed from enhancement to suppression at high *α* (Fig. 7A). Due to E-to-E STD, the effective excitatory-to-excitatory coupling decreases, resulting in a stable E subnetwork, and the network eventually becomes a non-inhibition-stabilized network (non-ISN) as demonstrated by the leading eigenvalues of the E subnetwork and E-VIP subnetwork switching from positive to negative with increasing bottom-up input (Fig. 7B). Interestingly, the response reversal of SST from enhancement to suppression does not occur at the same time as the network transitions from ISN to non-ISN. We proved that being an ISN is a necessary but not a sufficient condition to generate enhanced SST activity induced by top-down modulation, and non-ISNs cannot generate enhanced SST activity induced by top-down modulation (see SI Text). Consistent with our previous results, in the presence of E-to-E STD, the paradoxical response of SST is also linked to the change in SST activity induced by top-down modulation (Fig. 7C). More specifically, the network exhibits (no) paradoxical response of SST when top-down modulation increases (decreases) SST activity (Fig. 7C).

**Fig. 7.**
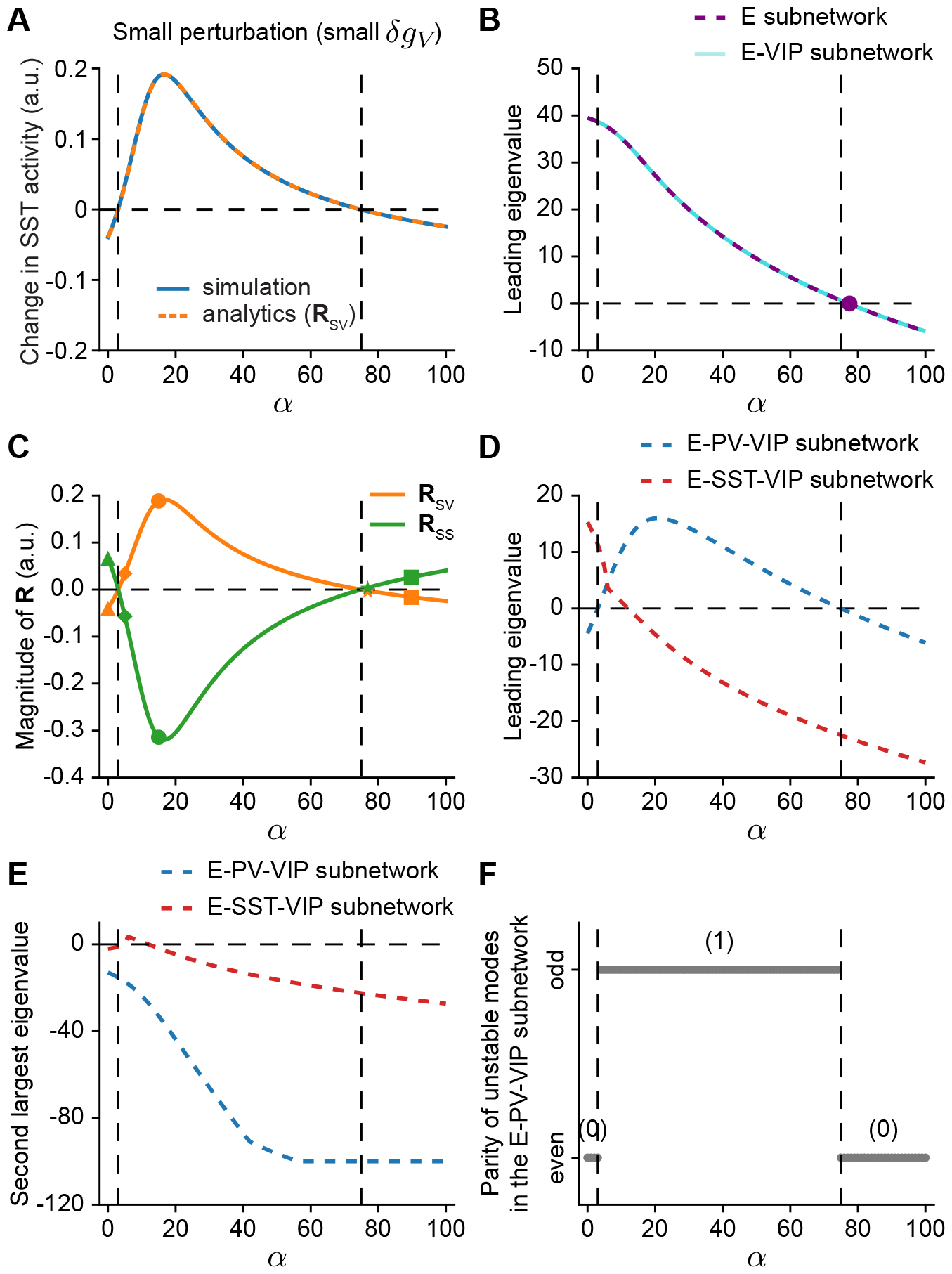
Modeling results are robust in the presence of E-to-E STD. (**A**) Change in SST activity as a function of bottom-up input *α* for networks also including E-to-E STD, showing numerical results and analytical predictions. The response reversal boundaries extracted from analytical calculations (**R**_*SV*_ = 0) are indicated by the vertical dashed lines. Here, *δg*_*V*_ = 0.1. (**B**) Leading eigenvalue of the E subnetwork and E-VIP subnetwork as a function of bottom-up input *α*. The leading eigenvalue eventually turns negative, indicating that the network becomes non-inhibition stabilized. The dot represents the *α* level at which the leading eigenvalues are zero. (**C**) Relationship between response reversal and the paradoxical response of SST. Analytical predictions of the change in SST response induced by an excitatory perturbation (*δg*_*S*_) to SST, **R**_*SS*_, and change in SST response induced by an excitatory perturbation (*δg*_*V*_) to VIP, **R**_*SV*_, as a function of bottom-up input *α*. Here, *δg*_*V*_ = *δg*_*S*_ = 0.1. (**D**) Similar to B but for the E-PV-VIP subnetwork and the E-SST-VIP subnetwork. (**E**) Similar to D but for the second largest eigenvalues. (**F**) Parity of the number of unstable modes in the E-PV-VIP subnetwork as a function of bottom-up input *α*. Numbers in the brackets indicate the amount of unstable modes.

Different from networks without E-to-E STD, by examining the leading eigenvalue of the E-PV-VIP and E-SST-VIP subnetwork (Fig. 7D), we observed a repertoire of interneuron-specific stabilization regimes and some novel regime transitions (Fig. 8). For instance, we observed a transition from being stabilized by PV but not SST (as reflected by a leading positive eigenvalue of the E-SST-VIP subnetwork and a leading negative eigenvalue of the E-PV-VIP subnetwork) to being stabilized by both PV and SST (as reflected by leading positive eigenvalues of the E-PV-VIP and E-SST-VIP subnetwork). We also observed a transition from being stabilized by SST but not PV (as reflected by a leading positive eigenvalue of the E-PV-VIP subnetwork and a leading negative eigenvalue of the E-SST-VIP subnetwork) to being stabilized by either PV or SST (as reflected by leading negative eigenvalues of the E-PV-VIP and E-SST-VIP subnetwork) (Fig. 7D, 8). We further confirmed these distinct regimes by injecting a transient excitatory perturbation into the excitatory population while clamping the activity of PV, or SST, or both PV and SST (Fig. S8). Despite novel regimes observed in the presence of E-to-E STD, the link between response reversal, paradoxical effects of SST, and the parity of unstable modes in the E-PV-VIP subnetwork remains unchanged (Fig. 7C-F). When top-down modulation decreases SST activity, the network exhibits no paradoxical response of SST, and the E-PV-VIP subnetwork has an even number of unstable modes. When top-down modulation increases SST activity, the network exhibits a paradoxical response of SST, and the E-PV-VIP subnetwork has an odd number of unstable modes implying that SST is required for network stabilization (Fig. 8).

**Fig. 8.**
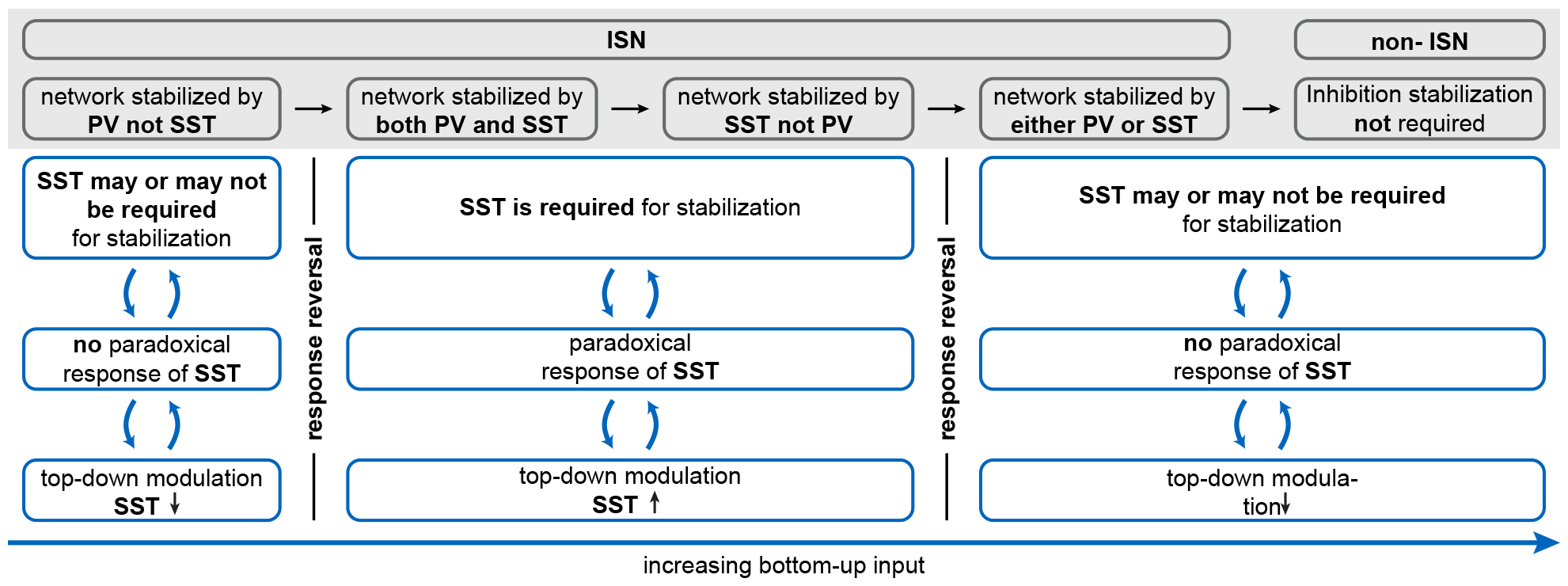
The relationship between response reversal, inhibition stabilization, and paradoxical response in networks also including E-to-E STD. At the low bottom-up input, top-down modulation decreases SST activity, and the network does not exhibit a paradoxical response of SST; thus, SST may not be required for stabilization, or SST may be required for stabilization, but the E-PV-VIP subnetwork has an even number of unstable modes. As demonstrated in Fig. 7, in this regime, the network is inhibition stabilized and stabilized by PV but not SST. With increasing bottom-up input, the response of SST induced by top-down modulation is reversed from suppression to enhancement. The network further exhibits the paradoxical effect in the response of SST and requires SST for stabilization with an odd number of unstable modes in the E-PV-VIP subnetwork. As demonstrated in Fig. 7, in this regime, the network is inhibition stabilized and stabilized by both PV of SST and then transitions into being stabilized by SST but not PV. Further increasing bottom-up input, the response of SST induced by top-down modulation is reversed from enhancement to suppression, and the network does not exhibit a paradoxical response of SST: thus, SST may not be required for stabilization, or SST may be required for stabilization, but the E-PV-VIP subnetwork has an even number of unstable modes. As demonstrated in Fig. 7, in this regime, the network is inhibition stabilized and stabilized by either PV or SST and finally transitions into a non-ISN. Note that while the relationship between response reversal, paradoxical effects, and inhibition stabilization marked in blue boxes does not depend on the choice of parameters, the possible interneuron-specific stabilization regimes shaded in gray are contingent on specific parameters.

In the presence of E-to-SST STF, the analytical expression of **K**_SV_ (Eq. 10), dictating the sign of the change of SST induced by top-down modulation, remains unchanged. It does not contain any E-to-SST dependent terms, suggesting that the emergence of response reversal is unaffected by E-to-SST STF (Fig. S9A). Consistent with the analysis, our simulation results show that adding E-to-SST STF does not alter the dynamics and the generation of response reversal (Fig. S9B, C). Given the omnipresence of short-term plasticity mechanisms in the mouse visual cortex amongst various populations and the centrality of SST to response reversal, we further incorporated short-term plasticity mechanisms in all connections considered in our model. These simulations show that response reversal can still be observed (Fig. S10). In addition, we examined how different inputs and network connectivity affect our results and found that response reversal is preserved in networks with varying inputs and connectivity strengths (Figs. S11, S12).

In conclusion, through multiple sensitivity analyses, we demonstrated the robustness of our findings to variations in short-term plasticity mechanisms, inputs, and network connectivity.

## Discussion

In this paper, we investigated how experimentally measured inhibitory short-term plasticity (iSTP) mechanisms enable model networks with one excitatory and three types of interneuron populations to perform a nonlinear computation known as response reversal. Using analytical calculations and numerical simulations, we identified that PV-to-E short-term depression (STD) is the iSTP mechanism critical for generating response reversal. We further clarified the relationship between response reversal, the paradoxical response of SST, and the interneuron-specific stabilization property of the network, making important links between well-known operating regimes of cortical network dynamics.

We made several assumptions that enabled us to analytically understand response reversal. First, we studied responses in the presence of bottom-up and top-down inputs relative to a baseline state, assuming that the network activity has reached a fixed point, and we did not consider scenarios like multistability (Hertäg and Sprekeler, 2019; Pietras et al., 2022) or oscillations (Veit et al., 2022). While multistability and oscillations have been observed in the brain (Wang, 2001; Buzsáki and Draguhn, 2004), the single stable fixed point assumed here is considered to be a realistic approximation of the awake sensory cortex (Miller, 2016).

Concerning the modeled short-term plasticity mechanisms, our analysis primarily focused on PV-to-E STD, PV-to-PV STD, PV-to-VIP STD, and SST-to-VIP STF. Additional simulations of networks also including E-to-E STD, or E-to-SST STF, or STP mechanisms in all existing synapses demonstrated the robust occurrence of response reversal in SST. In addition to the incorporated short-term plasticity mechanisms, substantial PV-to-SST STD has also been reported (Campagnola et al., 2022). However, experimental studies demonstrated negligible inhibition from PV to SST (Pfeffer et al., 2013), and hence we did not consider PV-to-SST STD.

Furthermore, our work models the neural input-output function as a rectified linear function, a characteristic feature of tightly balanced networks (van Vreeswijk and Sompolinsky, 1996, 1998). Without iSTP, our model network behaves like a linear network when all populations have positive activity. In addition to iSTP proposed in our study, several other factors can induce nonlinearities in the population response and, therefore, could contribute to the studied response reversal. Recent studies have suggested that cortical networks may operate in a loosely balanced regime, resulting in a supralinear input-output function (Ahmadian et al., 2013; Hennequin et al., 2018; Ahmadian and Miller, 2021; Ekelmans et al., 2023). Response reversal can also be generated by such a nonlinear input-output function (Garcia del Molino et al., 2017).

Finally, we modeled neurons of the same type as a homogeneous population governed by the same dynamics. In contrast, even within the same cell type, biological neurons have highly heterogeneous time constants and firing thresholds (Allen Institute for Brain Science, 2019; Cembrowski and Spruston, 2019). Such heterogeneity can theoretically also give rise to nonlinear population responses (Landau et al., 2016; Vegué and Roxin, 2019). Moreover, biological neurons possess complex morphologies (Jiang et al., 2015; Peng et al., 2021) and manifest nonlinear dendritic integrations (Poirazi et al., 2003; London and Häusser, 2005; Larkum et al., 2009; Tzilivaki et al., 2019). This suggests that the complete set of underlying mechanisms behind response reversal can be even richer and remains to be examined experimentally.

Our study makes several predictions. First, during top-down modulation, along with decreased SST activity, we also observed that the inhibition of the excitatory population increased at the steady state. Top-down modulation via VIP induces transient disinhibition, facilitating the growth of recurrent excitation and resulting in increased excitatory activity. This increased recurrent excitation is balanced by the concurrent increase in inhibition, which is a characteristic of inhibition-stabilized networks. This prediction can be tested experimentally by measuring excitatory and inhibitory currents to the excitatory neurons during top-down modulation.

Second, due to iSTP, locomotion-induced top-down modulation via VIP can reversely regulate SST response under different stimulus conditions. Although PV-to-E STD does not directly affect SST and VIP activity, surprisingly, our analysis suggests that PV-to-E STD is the determining mechanism underlying the generation of response reversal. Theoretical studies have demonstrated that inhibitory-to-inhibitory connections have the dominant impact on cortical dynamics, memory capacity, and working memory maintenance (Mongillo et al., 2018; Kim and Sejnowski, 2021). Here, our work suggests that the dynamics of inhibitory to excitatory synapses can be more important than those of inhibitory to inhibitory synapses to generate certain nonlinear phenomena.

Third, our theory reveals a correspondence between response reversal and a paradoxical response of SST in the presence of iSTP. More specifically, when the bottom-up stimulation condition switches from darkness to visual input, the impact of locomotion-induced top-down modulation via VIP on SST activity changes from suppression to enhancement. Once SST activity induced by top-down modulation gets elevated, the network exhibits a paradoxical response of SST. This correspondence can therefore be tested directly in future optogenetic experiments to see whether injecting excitatory (inhibitory) currents into the SST population indeed decreases (increases) its activity.

Fourth, our analysis shows that response reversal is tightly linked to the indispensability of SST for network stabilization. In darkness, when top-down modulation decreases SST activity, SST may not be required for network stabilization (i.e., solely PV can stabilize the network). In contrast, in the presence of visual stimuli, top-down modulation increases SST activity, and the network stabilization requires SST, and the E-PV-VIP subnetwork has an odd number of unstable modes. It is worth noting that when the network requires SST for stabilization, the network can require only SST but not PV (Fig. 5, S8) or both PV and SST for stabilization (Fig. S8). The novel observation that the network requires both PV and SST for stabilization is interesting and will require further investigation. Consistent with recent studies (Sanzeni et al., 2020), the network is inhibition-stabilized in all bottom-up stimulation conditions, even in darkness without visual stimuli. However, contrary to recent studies suggesting that PV is positioned to stabilize network activity (Bos et al., 2020), our work suggests that the specific inhibitory cell type stabilizing the network can change dynamically depending on the stimulation condition. As SST primarily targets dendrites of excitatory neurons (Kubota, 2014; Tremblay et al., 2016), stabilization through SST can be mechanistically realized via establishing a spatially precise E/I balance within individual dendritic segments. Recent experimental observations support the existence of such localized E/I balance at the dendritic segment level (Iascone et al., 2020), and SST is ideal for establishing the dendritic E/I balance and thus can provide an important source of stabilization. Furthermore, our results suggest a shift in inhibition source from PV to SST, typically accompanied by the occurrence of response reversal. Exploring the computational implications of this interneuron-specific inhibition shift and response reversal raises intriguing questions. Since PV neurons preferentially target perisomatic regions of excitatory neurons, whereas SST neurons target distal dendritic regions of excitatory neurons, the switch of dominant inhibition from soma to dendrite might prioritize inputs to perisomatic regions over inputs to distal dendritic regions and thus could be important for gating information (Udakis et al., 2020). Furthermore, inhibition also plays an important role in controlling plasticity (Letzkus et al., 2015). The redistribution of inhibition sources might imply different abilities of distinct interneurons to control plasticity in different regimes.

Last, for the network connectivity and the set of short-term plasticity mechanisms considered here, our results show that when SST is required for network stabilization and the E-PV-VIP subnetwork has an odd number of unstable modes, the network exhibits a paradoxical response of SST. Several studies have investigated the relationship between inhibition stabilization and the paradoxical effect in networks with multiple interneuron subtypes (Litwin-Kumar et al., 2016; Mahrach et al., 2020; Richter and Gjorgjieva, 2022; Palmigiano et al., 2023), in networks with short-term plasticity while ignoring different cell types (Sanzeni et al., 2020; Wu and Zenke, 2021), as well as in networks with multiple interneuron subtypes and short-term plasticity while ignoring cell type specificity (Wu and Gjorgjieva, 2023). How paradoxical effects of a given cell type relate to the number of unstable modes in subnetworks excluding that cell type has been studied in networks without short-term plasticity in a recent theoretical study (Miller and Palmigiano, 2020). Here, we revealed the relationship between interneuron-specific stabilization and the paradoxical effect in networks with multiple interneuron subtypes in the presence of a set of short-term plasticity mechanisms.

Taken together, our work sheds light on how experimentally identified iSTP mechanisms can generate response reversal, reveals the roles of individual iSTP mechanisms in response reversal, and uncovers the relationship between response reversal, the paradoxical effect, and interneuron-specific stabilization properties.

## Methods

### Response matrix

To investigate how the input to one particular population affects the response of any given population in the presence of short-term plasticity, we developed a general theoretical framework using linear perturbation theory. Using the separation of time scales for the rate dynamics and the short-term plasticity dynamics, we can write the system of equations introduced before (Eqs. 1 to 4) in matrix form while replacing the short-term plasticity variables with their steady-state values, as follows:

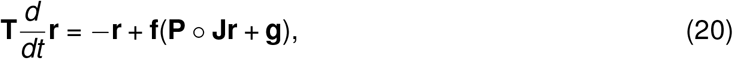

where **T** is a diagonal matrix of time constants of the firing rate dynamics, **r** a vector of firing rates of different populations, **f**(**x**) a vector of the rectified linear input-output function of the respective populations, **P** a matrix of the short-term plasticity variables, **J** the connectivity matrix, and **g** a vector of inputs to different populations. º denotes the element-wise product. The steady states of the short-term plasticity variables are obtained by setting the Eqs. 5 to 8 to 0. Note that since the steady states of the short-term plasticity variables are determined by the presynaptic activity, 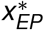, 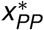, and 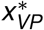are the same. If short-term plasticity is not present on the synapses from *j* to *i*, the corresponding element **P**_*ij*_ is 1 (for further details, see SI Text). By linearizing about the fixed point and ignoring higher-order terms, we obtain the following equation:

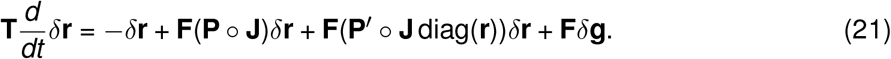

Here, *δ***r** is a vector containing the deviations of firing rates from their fixed point values. **F** is a diagonal matrix containing the derivatives of the input-output functions evaluated at the fixed point. **P**^′^ is a matrix containing the derivative of the short-term plasticity variables with respect to the corresponding presynaptic firing rate, evaluated at the fixed point. diag(**r**) is a diagonal matrix containing the firing rates of different populations. And *δ***g** is a vector containing the changes/perturbations of external inputs to different populations.

The fixed point solution of Eq. 21 quantifies the change in population rates *δ***r** to an input perturbation *δ***g**:

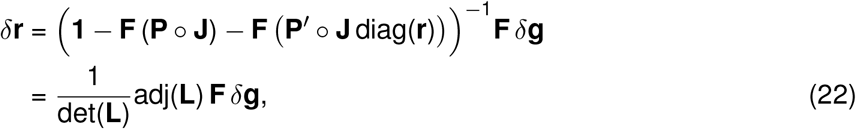

with

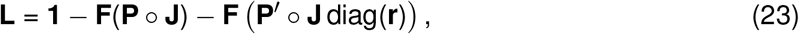

where **1** denotes the identity matrix, and ‘det’ and ‘adj’ represent the matrix’s determinant and adjugate, respectively.

By replacing *δ***g** with a diagonal matrix *δ***G** whose diagonal elements are *δ***g**, we can obtain a response matrix **R** as follows:

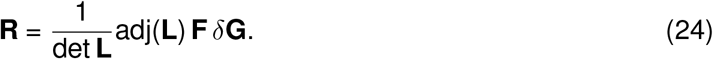

Importantly, the element **R**_*ij*_ provides the change in the steady-state rate response of population *I* caused by an input perturbation *δ***G**_*jj*_ to population *j*.

We further define a scalar *D* and a response factor matrix **K** as follows:

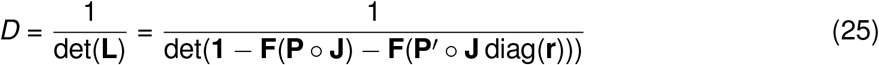

and

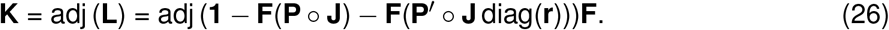

Then the response matrix **R** can be expressed as:

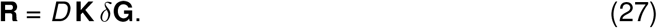

Note that if the network is stable, all eigenvalues of the Jacobian **T**^−1^(−**1**+**F**(**P** º **J**)+**F**(**P**^′^ º **J** diag(**r**))) have negative real parts. Therefore, all eigenvalues of **L** have positive real parts, and *D* is always positive (Fig. S4A).

To investigate how top-down modulation via VIP affects SST response in networks with iSTP, we can apply the theoretical framework introduced above and write the change of SST activity as a function of the change of the input to VIP. Since the derivatives of the rectified linear input-output functions are 1 in regimes where all cell populations have positive firing rates, we have

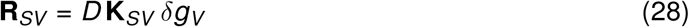

with

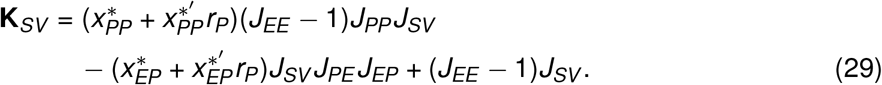

### Simulations

Simulations were performed in Python. All differential equations were implemented by Euler integration with a time step of 0.1 ms. The simulation duration was 9 seconds for each experiment. Top-down modulation was applied in the interval of 5 to 7 seconds. Networks were initialized using the parameters in the Supplementary Tables. Short-term plasticity variables were initially set to 1 and reached their steady-state values within the first second. Figures depict 6 seconds of network activity following 3 seconds of relaxation after initialization. Bottom-up input *α* was modeled in the interval [0, 20] with a step size of 0.5 unless stated otherwise. All simulation parameters are listed in the Supplementary Tables.

## Data Availability

The code used for model simulations is available on GitHub https://github.com/comp-neural-circuits/top-down-modulation-with-iSTP.

## Contributions

F.W., Y.K.W., and J.G. designed research; F.W. and Y.K.W. performed research; F.W. and Y.K.W. contributed new reagents/analytic tools; F.W. and Y.K.W. analyzed data; F.W., Y.K.W., and J.G. wrote the paper.

## Acknowledgments

We thank JaeAnn Dwulet, Elizabeth Herbert, Shreya Lakhera, and Fabio Veneto for commenting on the manuscript and the entire “Computation in Neural Circuits Group” for discussions. This work was supported by the European Research Council under the European Union’s Horizon 2020 research and innovation program (Grant Agreement No. 804824 to J.G.), the Deutsche Forschungsgemeinschaft in the Collaborative Research Centre 1080 (project C7 to J.G.), the Max Planck Society, and a grant from the Technical University of Munich (TUM Innovation Network Neurotech to J.G.). Y.K.W. is supported by the Add-on Fellowship of the Joachim Herz Foundation. We acknowledge the use of BioRender to generate Figure 1.

## Supporting Information

### Supporting Information Text

#### Response matrix

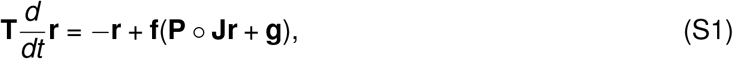

where **T** is a diagonal matrix of time constants of the firing rate dynamics, **r** a vector of firing rates of different populations, **f**(**x**) a vector of the rectified linear input-output function of the respective populations, **P** a matrix of the short-term plasticity variables, **J** the connectivity matrix, and **g** a vector of inputs to different populations. º denotes the element-wise product:

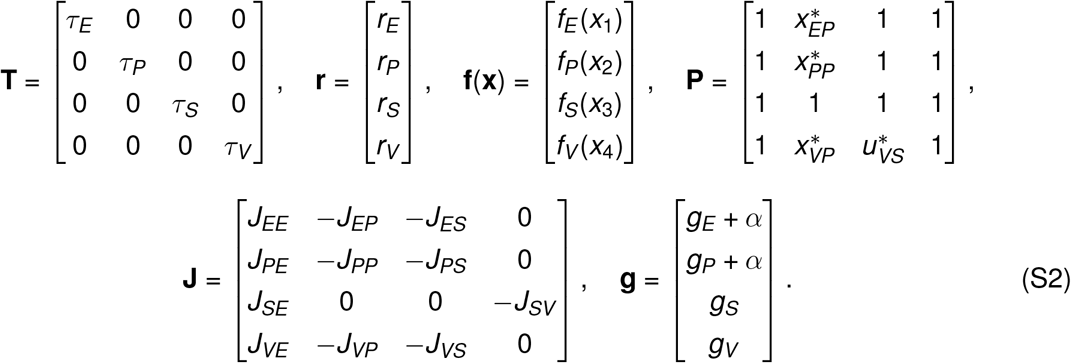

The steady states of the short-term plasticity variables are obtained by setting the Eqs. 5 to 8 to 0 and given by:

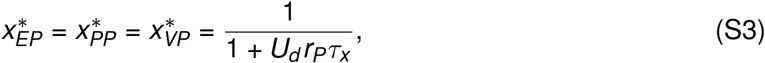

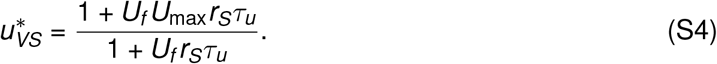

Note that since the steady states of the short-term plasticity variables are determined by the presynaptic activity, 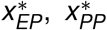, and 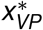are the same. If short-term plasticity is not present on the synapses from *j* to *i*, the corresponding element **P**_*ij*_ is 1.

By linearizing about the fixed point and ignoring higher-order terms, we obtain the following equation:

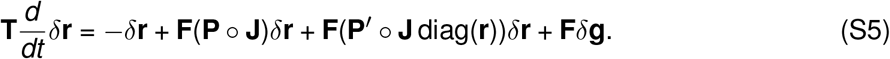

Here, *δ***r** is a vector containing the deviations of firing rates from their fixed point values. **F** is a diagonal matrix containing the derivatives of the input-output functions evaluated at the fixed point. **P**^′^ is a matrix containing the derivative of the short-term plasticity variables with respect to the corresponding presynaptic firing rate, evaluated at the fixed point. diag(**r**) is a diagonal matrix containing the firing rates of different populations. And *δ***g** is a vector containing the changes/perturbations of external inputs to different populations:

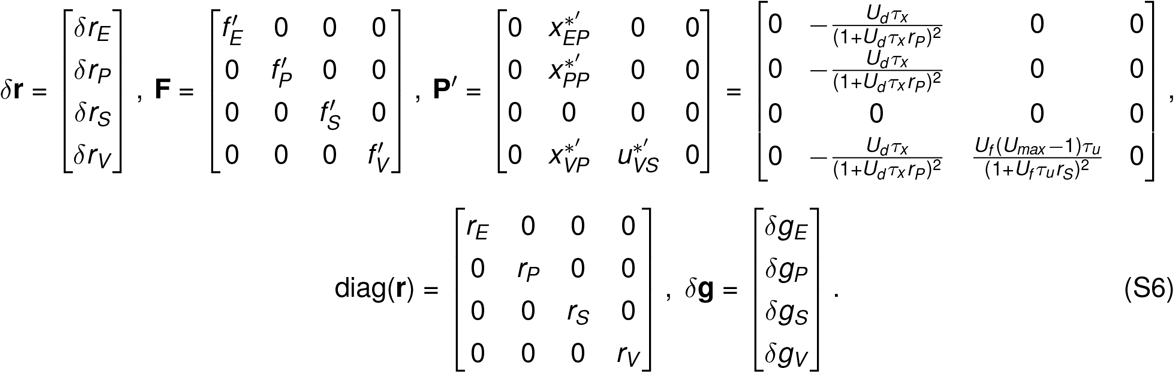

The fixed point solution of Eq. S5 quantifies the change in population rates *δ***r** to an input perturbation *δ***g**:

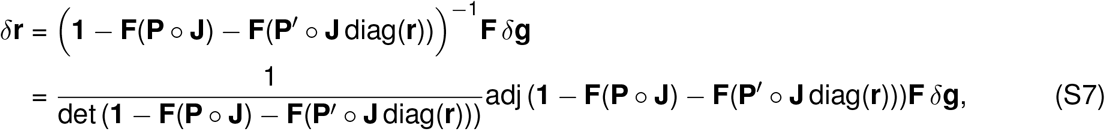

where **1** denotes the identity matrix, and ‘det’ and ‘adj’ represent the matrix’s determinant and adjugate, respectively. By replacing *δ***g** with a diagonal matrix *δ***G** whose diagonal elements are *δ***g**, we can obtain a response matrix **R** as follows:

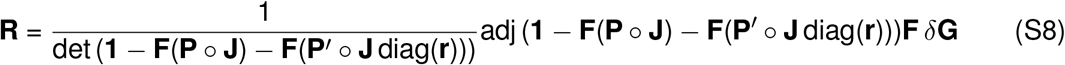

with

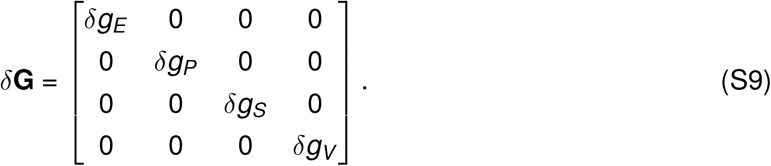

Importantly, the element **R**_*ij*_ provides the change in the steady-state rate response of population *i* caused by an input perturbation *δ***G**_*jj*_ to population *j*. We further define a scalar *D* and a response factor matrix **K** as follows:

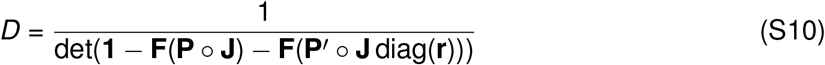

and

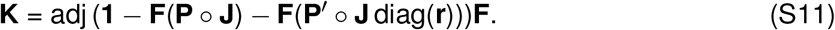

Then the response matrix **R** can be expressed as:

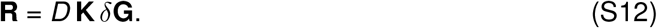

Note that if the network is stable, all eigenvalues of the Jacobian **T**^−1^(−**1**+**F**(**P** º **J**)+**F**(**P**^′^ º **J** diag(**r**))) have negative real parts. Therefore, all eigenvalues of **1** − **F**(**P** º **J**) − **F**(**P**^′^ º **J** diag(**r**)) have positive real parts and *D* is always positive (Fig. S4A).

To investigate how top-down modulation via VIP affects SST response in networks with iSTP, we can apply the theoretical framework introduced above and write the change of SST activity as a function of the change of the input to VIP:

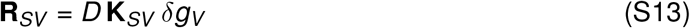

with

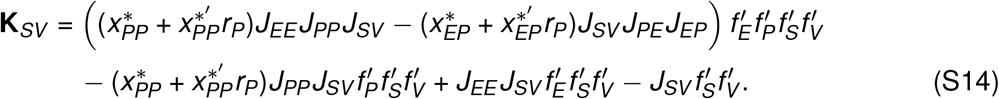

To have response reversal of SST from suppression to enhancement, we need the change of SST activity induced by top-down modulation via VIP to switch from negative to positive as bottom-up input increases. In other words, we need a sign change of **R**_*SV*_ from negative to positive as the bottom-up input increases. Since *D* and *δg*_*V*_ are positive, **K**_*SV*_ is the only term that can switch the sign of **R**_*SV*_.

In the regime where all cell populations have positive firing rates, the derivatives of the rectified linear input-output functions are 1. Therefore, **K**_*SV*_ can be simplified to:

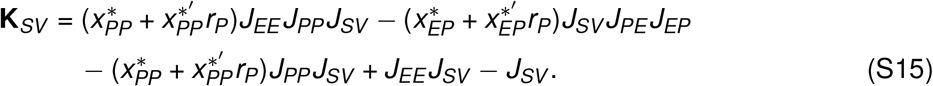

We can further separate **K**_*SV*_ into an STP-dependent part 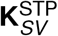 and a non-STP-dependent part 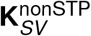:

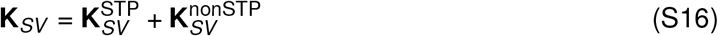

with

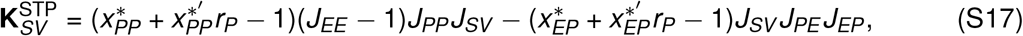

and

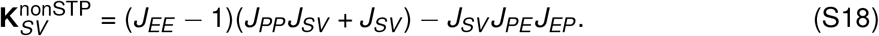

The STP-dependent part 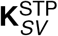 can be expressed as a sum of a PV-to-E STD-dependent term 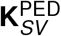 and a PV-to-PV STD-dependent term 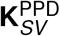:

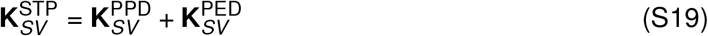

with

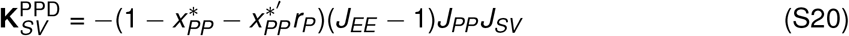

and

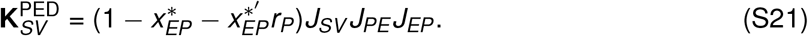

Furthermore,

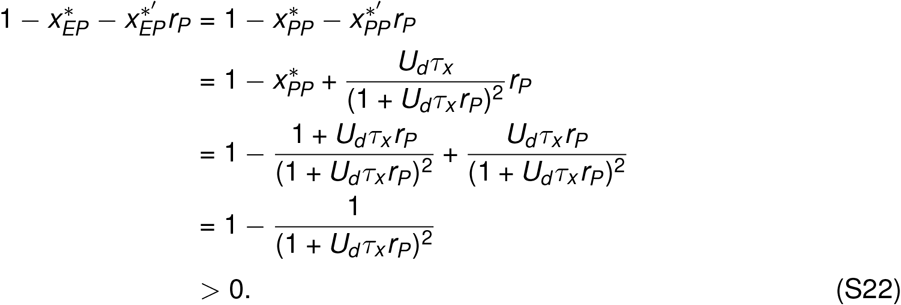

The PV-to-E STD-dependent term 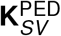, therefore, is always positive. Consistent with recent experimental studies (Sanzeni et al., 2020), the network is initialized in an inhibition-stabilized regime when the animal receives no stimulus in darkness. In other words, *J*_*EE*_ is greater than 1 (see section ‘Interneuron-specific stabilization property’), resulting in the PV-to-PV STD-dependent term 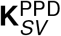 being always negative. Taking the derivative of Eq. S22 with respect to *r*_*P*_, we get:

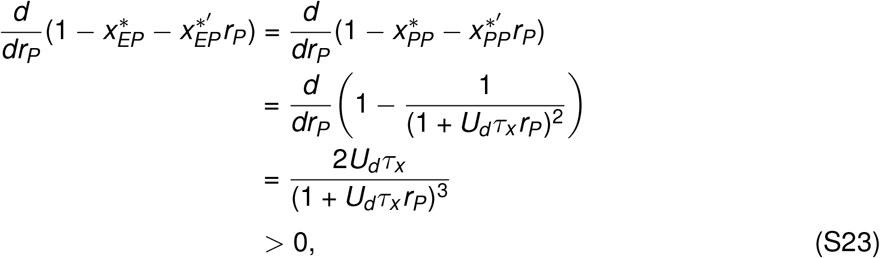

resulting in a continuous increase (decrease) in 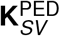 and a continuous decrease (increase) in 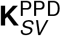with increasing (decreasing) rate of PV. Since PV activity *r*_*P*_ increases (decreases) with greater (smaller) bottom-up input *α* (Fig. S2), 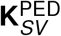 increases (decreases) while 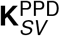 decreases (increases) (Fig. S4C). For response reversal of SST from negative to positive with increasing *α*, **K**_*SV*_ needs to switch from negative to positive. Therefore, 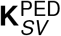, rather than 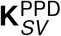, is the critical short-term plasticity term for response reversal from suppression to enhancement.

### Interneuron-specific stabilization property

Network stabilization can be determined by the leading eigenvalue of the Jacobian of the system. The Jacobian of our network with iSTP is given by:

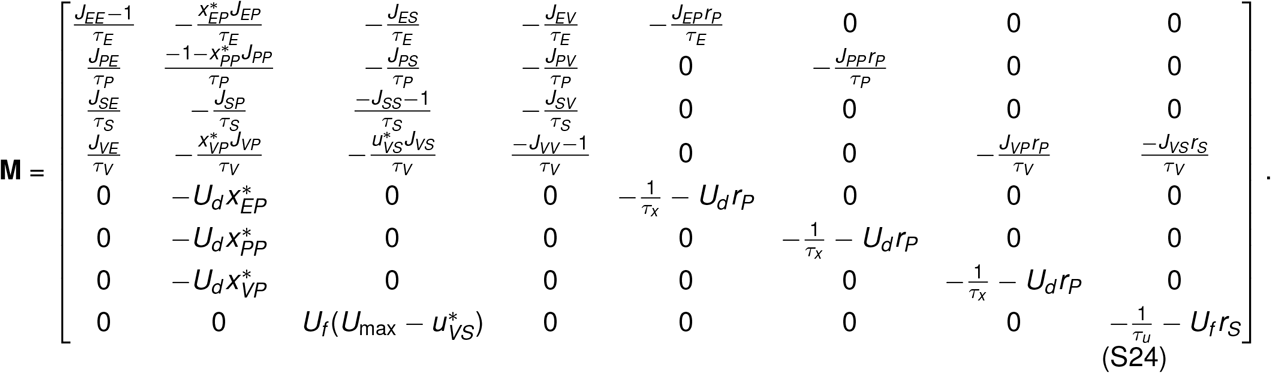

A positive leading eigenvalue implies that the fixed point is unstable. In other words, a transient perturbation to the system leads to a deviation from the original fixed point. In contrast, a negative leading eigenvalue implies that the fixed point is stable. Namely, the system will return to the original fixed point after transient perturbation.

To identify if the network is inhibition stabilized, we computed the leading eigenvalue of the Jacobian of the E subnetwork. In this case, all inhibitory populations and their related STP variables are omitted. The leading eigenvalue of the Jacobian of the E subnetwork is 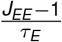. Therefore, when *J*_*EE*_ is larger than 1, the excitatory subnetwork is unstable, and the network is inhibition stabilized, i.e. in the ISN regime. In alignment with recent experimental studies showing that in darkness, when animals receive no stimulus, the network is inhibition stabilized (Sanzeni et al., 2020), we thus set *J*_*EE*_ to be larger than 1.

Since

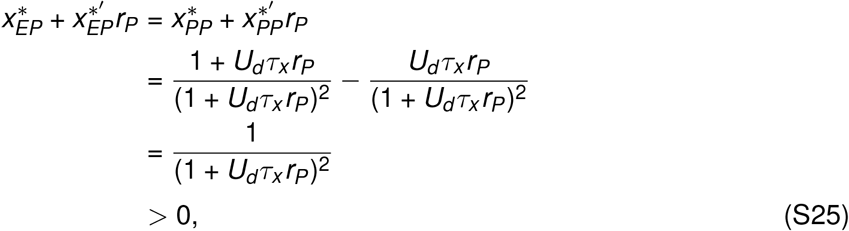

it is easy to see that **K**_*SV*_ (Eq. 29) is negative when *J*_*EE*_ − 1 *<* 0. This implies that for any network whose E subnetwork is stable (i.e., the entire network is not inhibition stabilized), top-down modulation via VIP decreases SST activity. The network, therefore, needs to be inhibition stabilized, i.e. the E subnetwork has to be unstable, to obtain an enhanced effect of SST response induced by top-down modulation.

To identify if the network is inhibition stabilized by a specific interneuron subtype, we compute the leading eigenvalue of the subnetwork in which a specific interneuron subtype and the corresponding STP mechanisms are omitted. If the leading eigenvalue is positive (negative), the subnetwork without this interneuron population is unstable (stable), suggesting the omitted interneuron sub-type is required (not required) for stabilization. This interneuron-specific stabilization property can also be probed by, at the same time, freezing the respective inhibitory population, transiently perturbing the excitatory population and observing potential changes in the fixed point dynamics.

To reveal the relationship between the response reversal of SST, the paradoxical effect, and interneuron-specific stabilization, we compute the Jacobian for the E-PV-VIP subnetwork in which the SST population and its related STP variable are omitted. The Jacobian of the E-PV-VIP sub-network is given by:

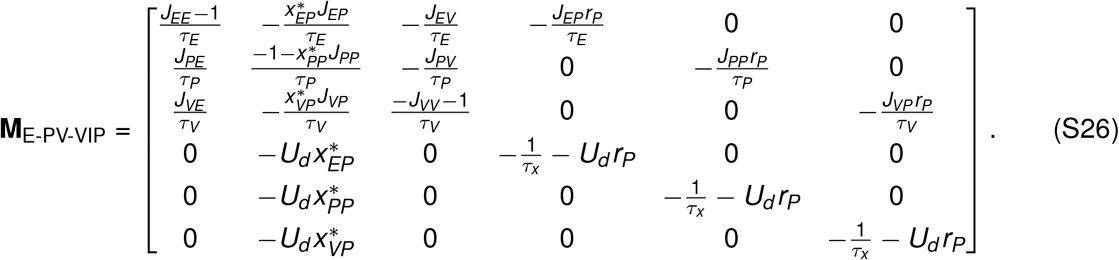

Therefore, the determinant of the Jacobian is given by

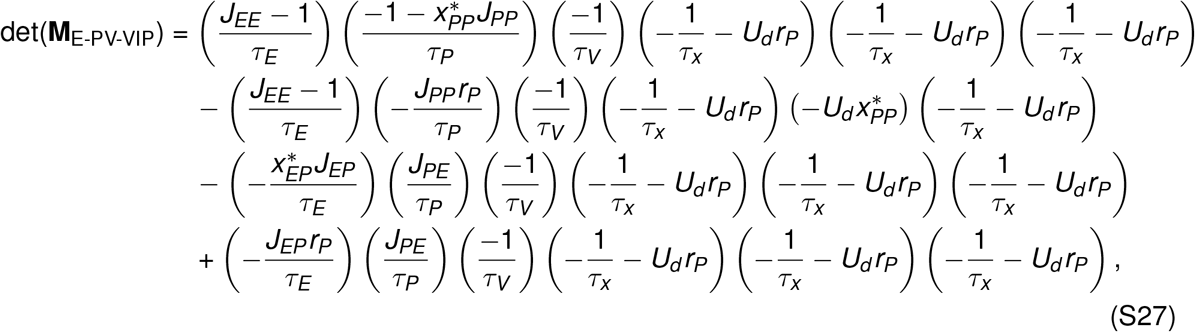

which becomes:

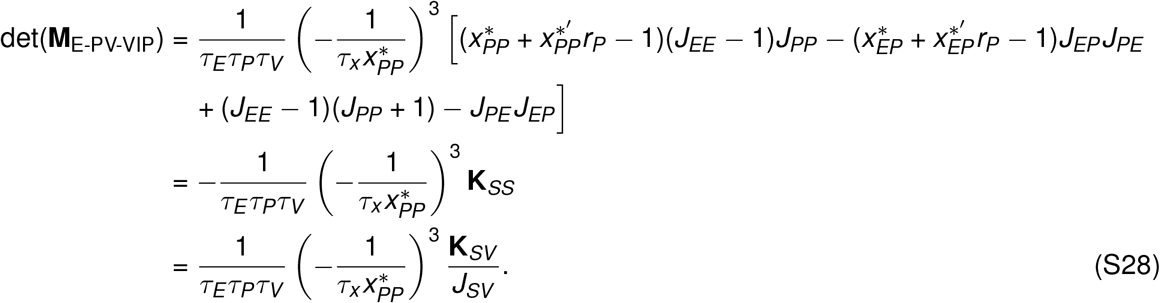

In the network we considered here, because of the short-term plasticity mechanisms, the Jacobian of the E-PV-VIP subnetwork is a 6-by-6 matrix. Since det(**M**_E-PV-VIP_) is the product of the eigenvalues of the Jacobian for the E-PV-VIP subnetwork, a negative det(**M**_E-PV-VIP_) implies an odd number of positive eigenvalues and consequently an odd number of unstable eigenvectors/modes in the E-PV-VIP subnetwork, suggesting that SST is required for network stabilization. However, the network may also require SST for stabilization if det(**M**_E-PV-VIP_) is positive, for instance, when the Jacobian of the E-PV-VIP subnetwork has an even number of unstable modes. At the same time, an odd number of unstable modes in the E-PV-VIP subnetwork also implies a negative **K**_*SS*_ and a positive **K**_*SV*_, suggesting that SST responds paradoxically, and top-down modulation via VIP increases SST activity. In contrast, an even number of unstable modes in the E-PV-VIP subnetwork implies a negative **K**_*SS*_ and a positive **K**_*SV*_.

Further, to investigate the relationship between PV stabilization and response reversal of SST, we compute the Jacobian for the E-SST-VIP subnetwork, which is given by:

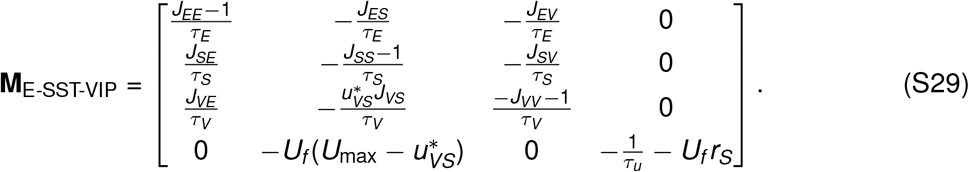

Its determinant is then given by

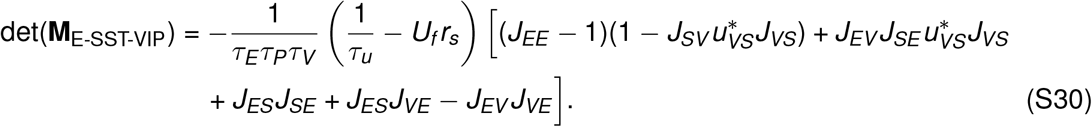

We found no relation between the determinant of the E-SST-VIP subnetwork det(**M**_E-SST-VIP_) and the response reversal condition **K**_*SV*_. The determinant can switch its sign independent of the response of SST and vice versa. This implies that the requirement of PV for network stabilization cannot be linked to the response reversal condition of SST. As a result, the PV stabilization property can change independently with different bottom-up inputs and is thus parameter-dependent.

### Networks also including E-to-E STD

In the presence of E-to-E STD, we have

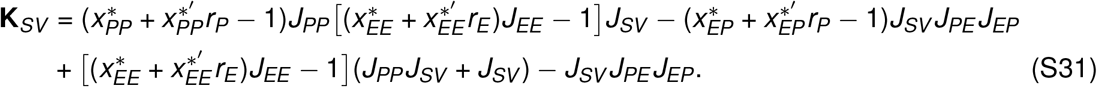

with

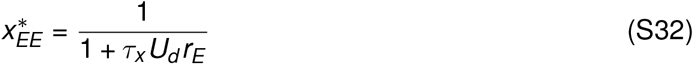

and

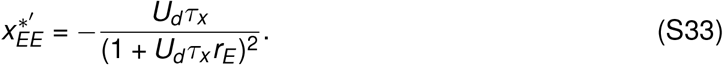

Furthermore, we have

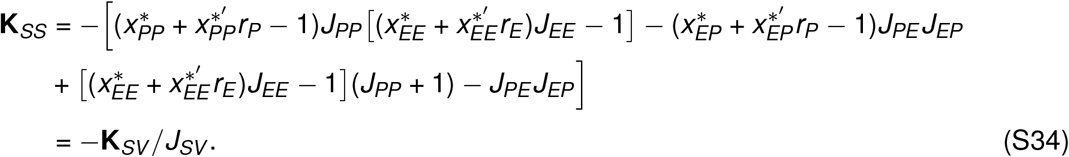

Note that in the presence of E-to-E STD, we still have

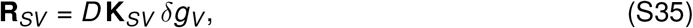

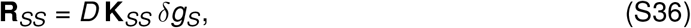

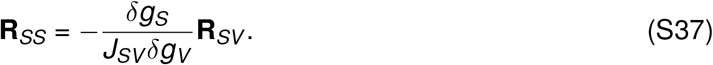

Despite the fact that E-to-E STD affects the amplitude of *D*, the relationship between **R**_*SV*_ and **R**_*SS*_ remains unchanged.

To investigate how **K**_*SV*_ relates to inhibition stabilization, we analyzed the Jacobian of the E-to-E subnetwork now including E-to-E STD, **M**_*EE*_, given by:

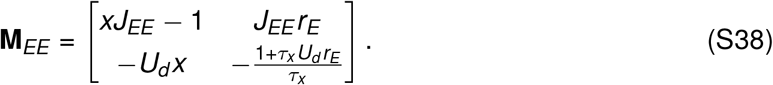

If the E subnetwork is stable (i.e., the entire network is not inhibition stabilized), the determinant of the Jacobian det(**M**_*EE*_) is positive, namely,

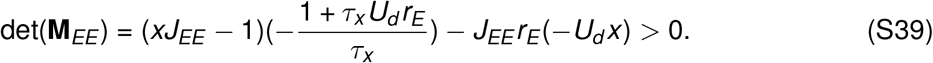

At the steady state, the above equation can be written as follows

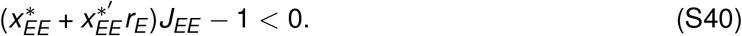

Since 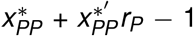 and 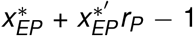 are positive (Eq. S22), it is easy to see that when 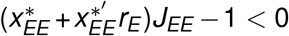(for instance, when the network is not inhibition stabilized), **K**_*SV*_ (Eq. S31) is always negative. Therefore, to have increased SST activity induced by top-down modulation (i.e., **K**_*SV*_ *>* 0), the network has to be inhibition stabilized.

Furthermore, as bottom-up input increases, *r*_*E*_ also increases. Since

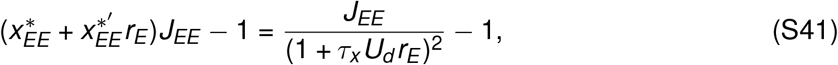

to generate elevated SST activity by top-down modulation at a high level of bottom-up input,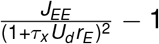 needs to be positive. As a result, *J*_*EE*_ has to be large and/or *τ*_*x*_ *U*_*d*_ has to be small. Note that as shown in Eq. (S32), greater *τ*_*x*_ *U*_*d*_ implies a stronger depression of E-to-E connection strength.

### The Response Reversal Index

In the exploration of the robustness of our findings under various perturbations, we conducted sensitivity analyses encompassing inputs, network connectivity, and short-term plasticity mechanisms. For a concise evaluation of the transition in SST response from suppression to enhancement induced by top-down modulation and the consequent emergence of response reversal, we introduce the Response Reversal Index (RRI) as follows:

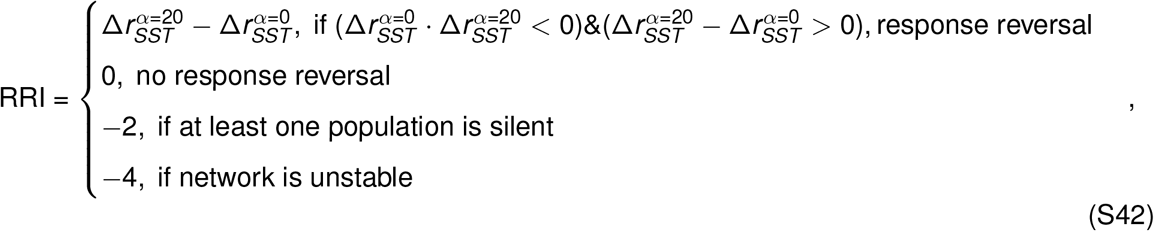

where 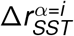 represents the change in SST response induced by top-down modulation at the bottom-up input *i*. Note that a positive RRI indicates a response reversal of SST from negative to positive.

### Sensitivity analysis to background inputs and connectivity matrix

We tested whether our results were sensitive to the choices of background inputs *g* and the ratio of *g* and top-down modulatory input *c*. We varied the mean background input 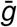 (i.e., the average background input to four different populations) by randomly and independently sampling the corresponding background inputs to individual populations from a uniform distribution (see Methods). We found that response reversal is observed at different levels of background input (Fig. S11A). Furthermore, we covaried the top-down modulatory input *c* and found that our results are robust for a wide range of ratios between background input and top-down modulatory input (Fig. S11B). These results suggest that our results remain robust across a wide range of bottom-up and top-down inputs.

Although our initial network connectivity is constrained by previous experimental studies (Pfeffer et al., 2013), strengths of individual synapses have huge variability. We therefore examined whether our results hold for a variety of networks with different connectivity strengths. We varied the excitatory to excitatory connectivity strength *J*_*EE*_ and found that non-inhibition stabilized networks (i.e., the E subnetwork is stable) are unable to generate response reversal (Fig. S12A).

We mathematically proved that the network needs to be inhibition stabilized (i.e. the E subnetwork has to be unstable) to obtain an enhanced effect on SST response induced by top-down modulation (see Methods). For various *J*_*EE*_, the inhibition-stabilized network can exhibit response reversal (Fig. S12A). Further increasing *J*_*EE*_ eventually leads to unstable networks, which can be prevented by increasing inhibitory weights. This also rescues response reversal (Fig. S12B). As our model suggests a shift in the primary source of inhibition from PV to SST when response reversal is observed, we examined how the ratio between *J*_*EP*_ and *J*_*ES*_ affects our conclusions. We found that a large fraction of the sampled inhibition-stabilized networks with a wide range of ratios between *J*_*EP*_ and *J*_*ES*_ are capable of generating response reversal (Fig. S12C, D). As the PV-to-E connection is subject to short-term depression, we then examined how the effective ratio between *J*_*EP*_ and *J*_*ES*_ affects response reversal. We, therefore, multiplied *J*_*EP*_ with the corresponding short-term plasticity variable *x*_*EP*_. Response reversal is observed for effective ratios larger than 1 as well as smaller than 1 (Fig. S12E). This is also the case for a wide range of connection strengths between E and PV (Fig. S12F), between E and SST (Fig. S12G), as well as *J*_*SE*_ and *J*_*PE*_ (Fig. S12H). These findings indicate the robustness of our results across inhibition-stabilized networks with varying connectivity strengths.

## Supplementary Figures

**Fig. S1.**
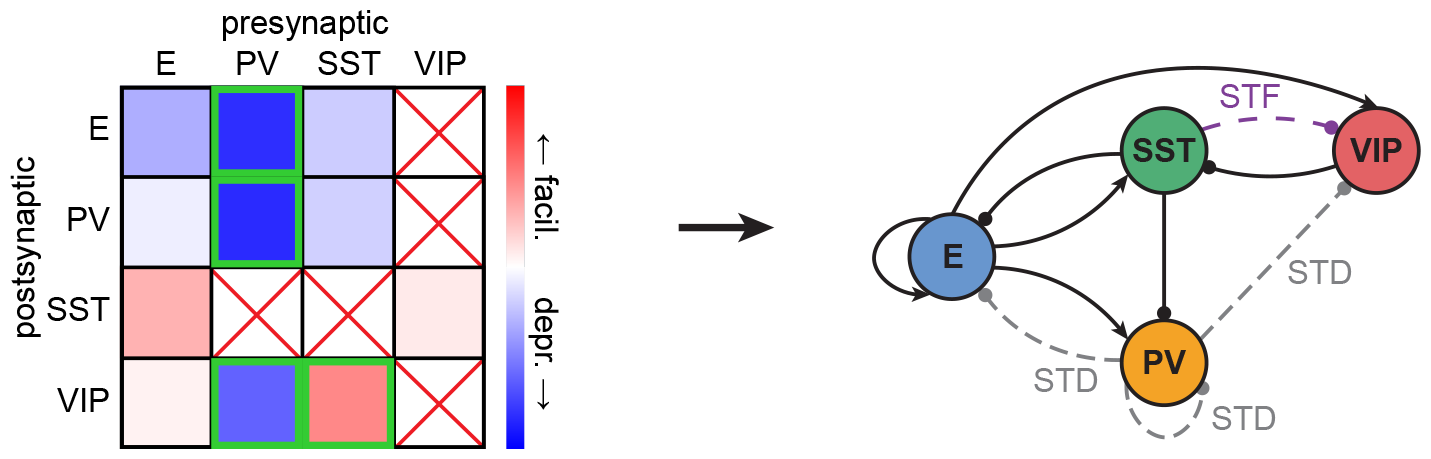
Short-term plasticity mechanisms included in the network model. Left: Different degrees of short-term facilitation (STF) and depression (STD) at different synapses measured by the Allen Institute (Campagnola et al., 2022). Green boxes indicate the four most pronounced mechanisms, the only ones included in the model. Red crosses denote weak connections, as reported in (Pfeffer et al., 2013). Therefore, not considered in the model. Right: Network schematic including the four most pronounced STD and STF mechanisms.

**Fig. S2.**
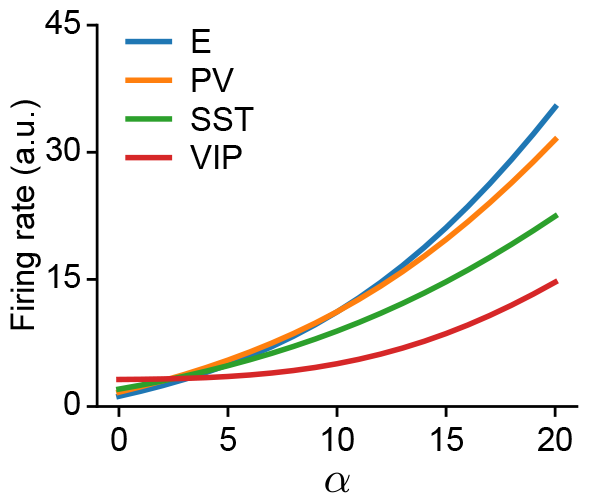
Network activity as a function of bottom-up input *α* to E and PV.

**Fig. S3.**
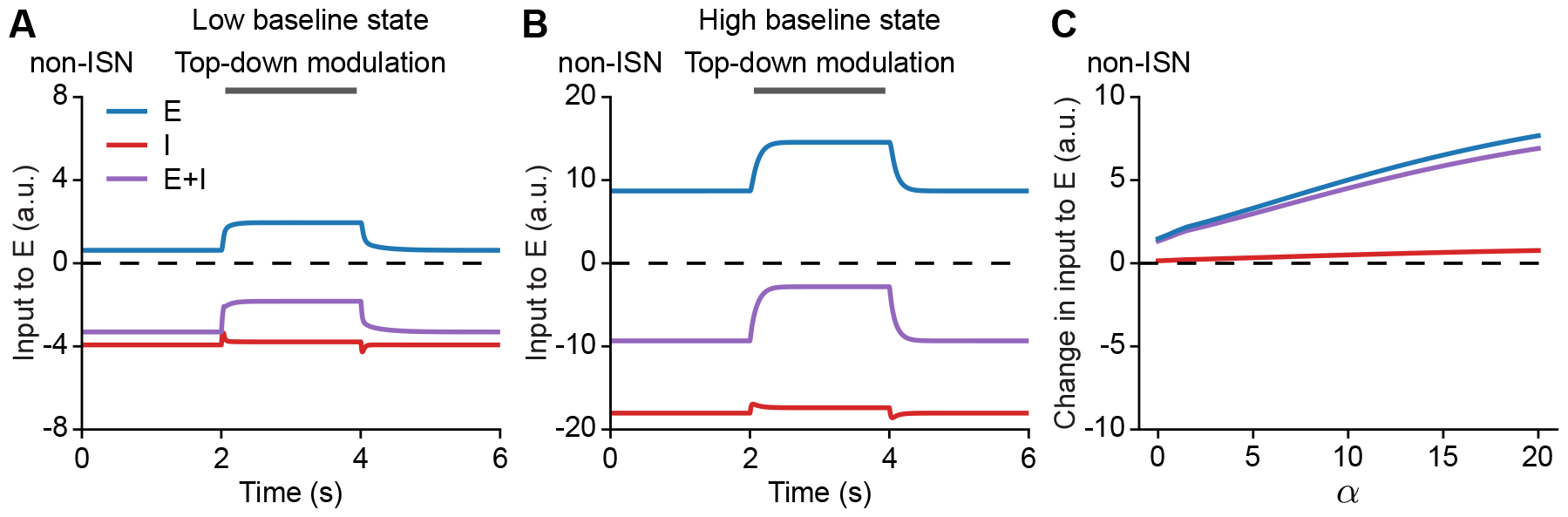
Non-inhibition-stabilized networks (non-ISNs) do not show elevated total inhibitory inputs to the excitatory population. (**A**) Input to the E population at *α* = 0 corresponding to darkness, i.e., no sensory stimulation. Top-down modulation via VIP is applied during the interval from 2 to 4 s (gray bar). Different colors indicate different sources: input from the E population, input from the I populations, and the sum of the inputs from the E and I populations. (**B**) Same as A but at *α* = 15 corresponding to sensory stimulation. (**C**) Change in different sources of recurrent inputs to the E population measured between baseline and at steady state during top-down modulation as a function of bottom-up input *α*.

**Fig. S4.**
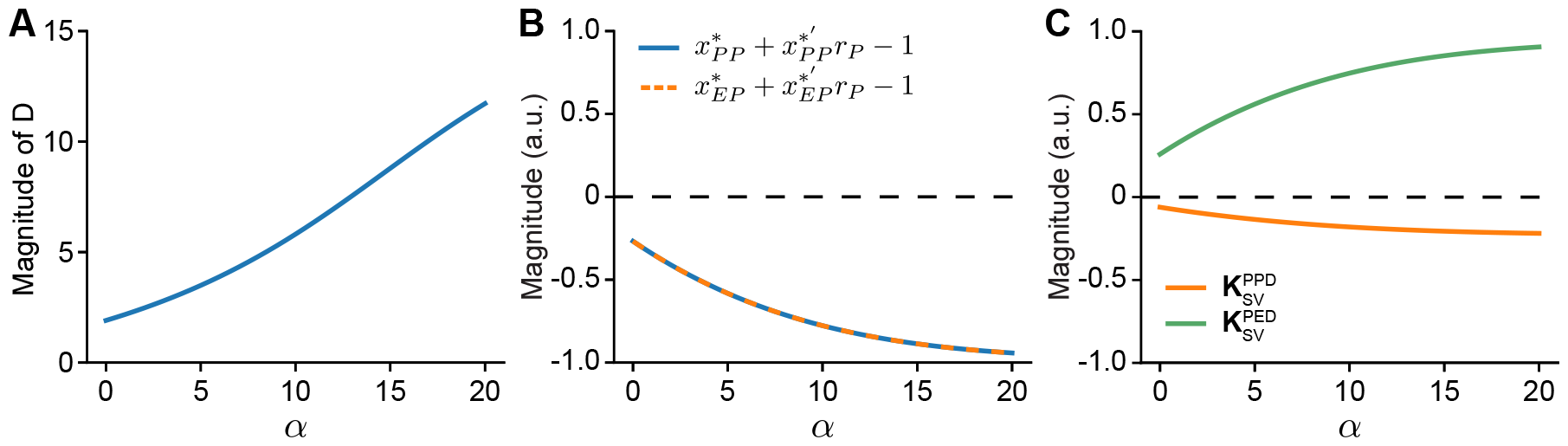
Bottom-up input changes the magnitude of individual components of **R**_*SV*_. (**A**) Magnitude of *D* in Eq. (S10) as a function of bottom-up input *α*. (**B**) Magnitude of the PV-to-PV STD-related term and the PV-to-E STD-related term (see Eqs. (S19)–(S21)) as a function of bottom-up input *α*. (**C**) PV-to-PV STD-dependent term 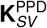 and the PV-to-E STD-dependent term 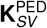 (see Eqs. (S19)–(S21)) as a function of bottom-up input *α*.

**Fig. S5.**
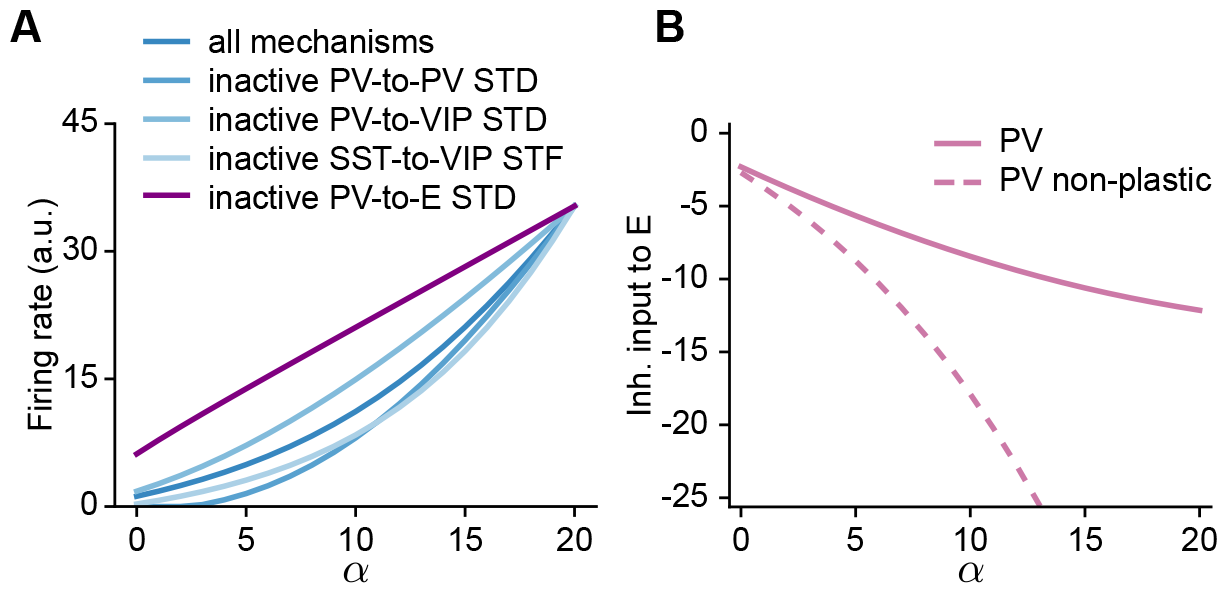
PV-to-E STD generates an effective supralinear input-output relation. (**A**) Excitatory activity as a function of bottom-up input *α* for different network configurations marked with different colors. (**B**) Inhibitory input from PV to E as a function of bottom-up input *α*. The solid line represents the input when PV-to-E STD is active, whereas the dashed line represents the input when PV-to-E STD is inactive.

**Fig. S6.**
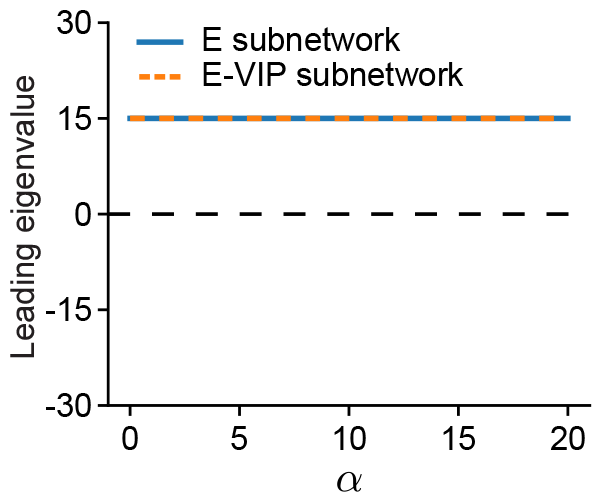
Leading eigenvalue of the E subnetwork and the E-VIP subnetwork subnetwork as a function of bottom-up input *α*.

**Fig. S7.**
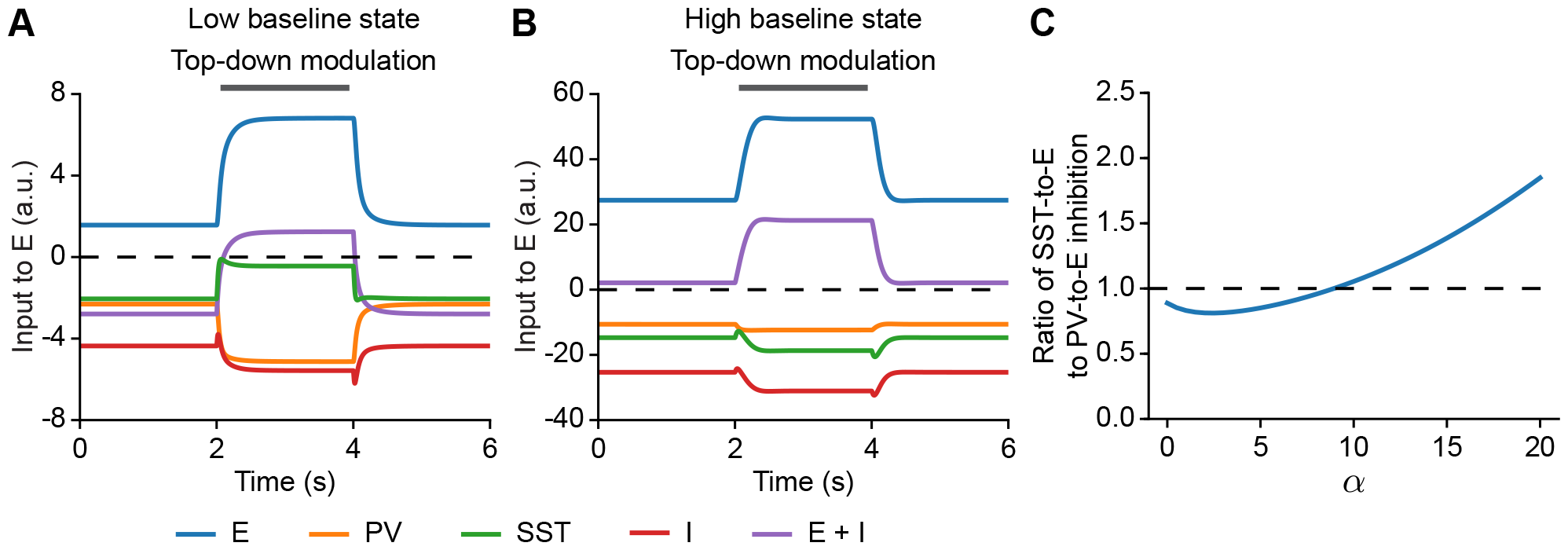
Bottom-up inputs shift the prevalence of inhibition received by the excitatory population (E) from PV to SST. (**A**) Input to the E population at *α* = 0 corresponding to darkness, i.e., without sensory stimulation. Top-down modulation via VIP is applied during the interval from 2 to 4 s (gray bar). Different colors indicate different sources: input from the E population, input from the PV population, input from the SST population (green), I populations, and the sum of the inputs from the E and I populations. (**B**) Same as A but at *α* = 15 corresponding to sensory stimulation. (**C**) The ratio of SST to PV inhibition received by the E population at baseline as a function of bottom-up input *α*.

**Fig. S8.**
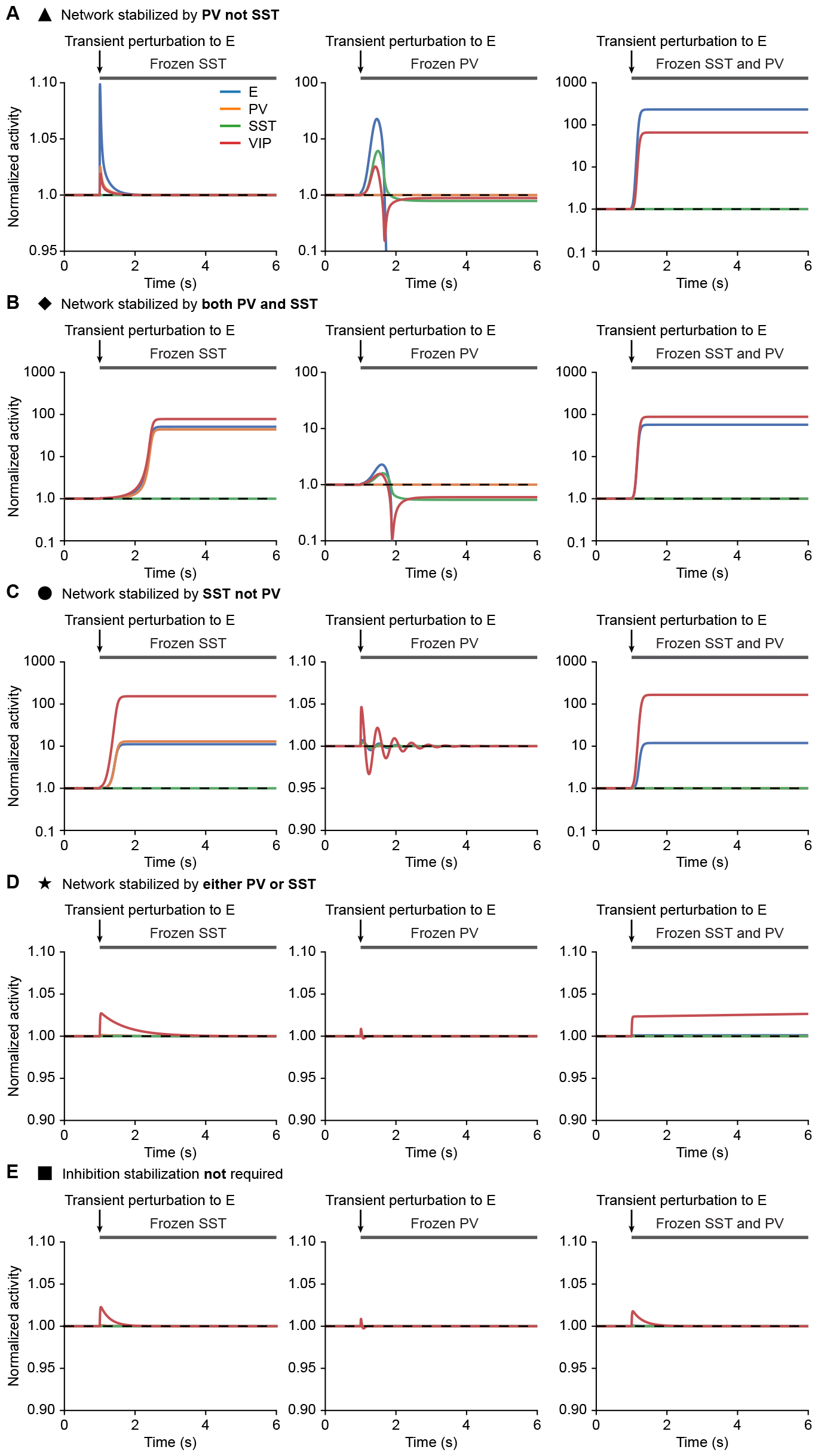
Responses of networks also including E-to-E STD to transient perturbations to the excitatory population while freezing corresponding inhibition at different baseline states, indicated in Fig. 7D. (**A**) Normalized activity when injecting additional excitatory current into E while freezing SST, or PV, or both, respectively, for a bottom-up input of *α* = 0. A small transient excitatory perturbation to the excitatory population is introduced at the time point marked with arrows. The periods in which inhibition is frozen are marked with the gray bar. The network is inhibition stabilized and stabilized by PV but not SST. Only freezing PV or both PV and SST causes a deviation from the original fixed point. (**B**) Similar to A but for a bottom-up input *α* = 5. The network is inhibition stabilized and requires SST and PV for stabilization, as a transient perturbation leads to a deviation from the original fixed point in all freezing experiments. (**C**) Similar to A but for a bottom-up input *α* = 15. The network is inhibition stabilized and stabilized by SST but not PV. Only freezing SST or both PV and SST causes a deviation from the original fixed point. (**D**) Similar to A but for a bottom-up input *α* = 77. The network is inhibition stabilized and stabilized by either PV or SST. Only freezing both PV and SST causes a deviation from the original fixed point. (**E**) Similar to A but for a bottom-up input *α* = 90. The network is no longer inhibition stabilized. Despite frozen inhibition, a transient perturbation does not result in a deviation from the original fixed point.

**Fig. S9.**
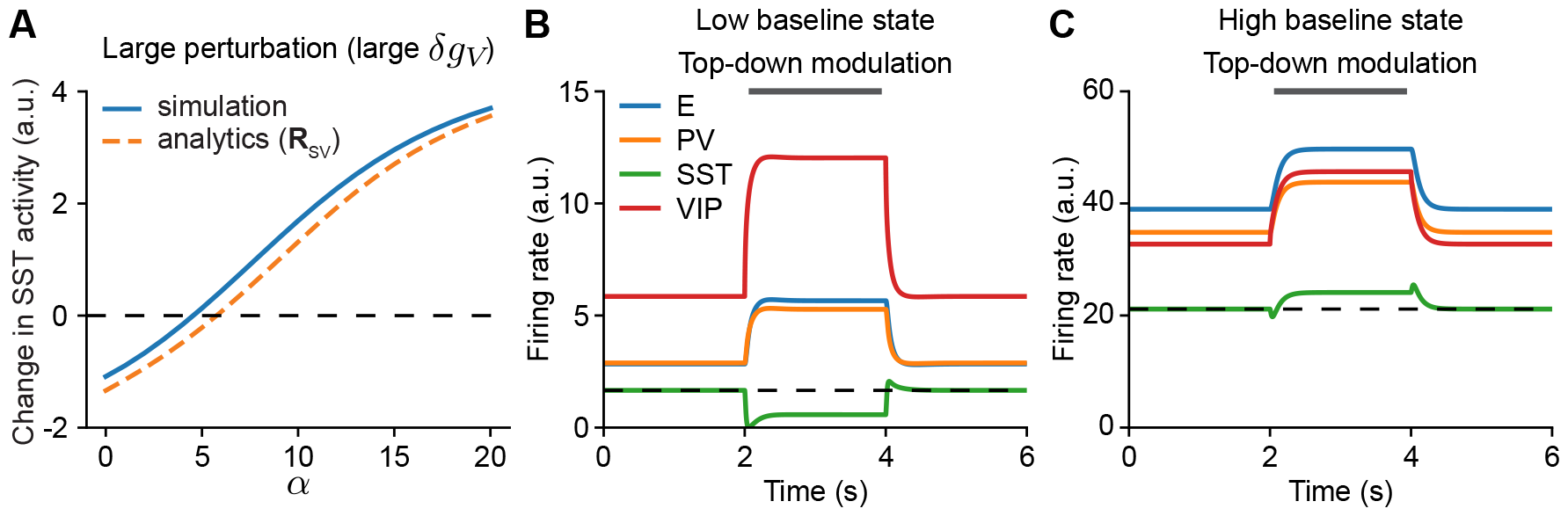
Response reversal in networks also including E-to-SST STF. (**A**) In networks also including E-to-SST STF, analytical prediction of the change in SST population response induced by the perturbation to VIP (**R**_*SV*_, Eq. 10) qualitatively match with numerical simulation for a large perturbation (*δg*_*V*_ = 3). (**B**) Network responses, including E-to-SST STF, to top-down modulation without any bottom-up input (*α* = 0) corresponding to darkness without sensory stimulation. Top-down modulation via VIP is applied during the interval from 2 to 4 s (gray bar). Different colors denote the activity of different populations. The dashed line represents the initial activity level of SST. (**C**) Similar to B but at *α* = 15 corresponding to sensory stimulation. Simulations were performed with top-down modulation *c* = 3 and 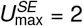.

**Fig. S10.**
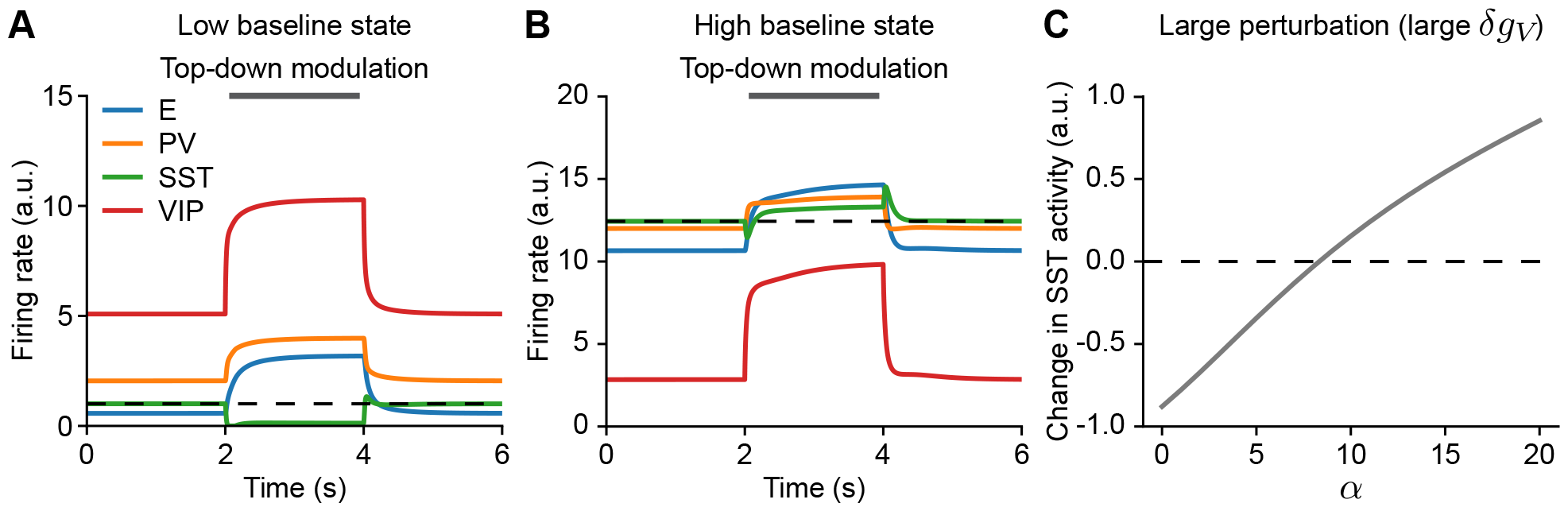
Networks with multiple additional short-term plasticity mechanisms are able to generate response reversal. (**A**) Network responses, including all short-term mechanisms observed in (Campagnola et al., 2022), on existing connections, to top-down modulation without any bottom-up input (*α* = 0) corresponding to darkness without sensory stimulation. Top-down modulation via VIP is applied during the interval from 2 to 4 s (gray bar). The dashed line represents the initial activity level of SST. (**B**) Same as A but at *α* = 15 corresponding to sensory stimulation. (**C**) Change in SST response induced by top-down modulation to VIP as a function of bottom-up input *α* in networks with short-term plasticity on all existing connections. Simulations were performed with top-down modulation *δg*_*V*_ = 3.

**Fig. S11.**
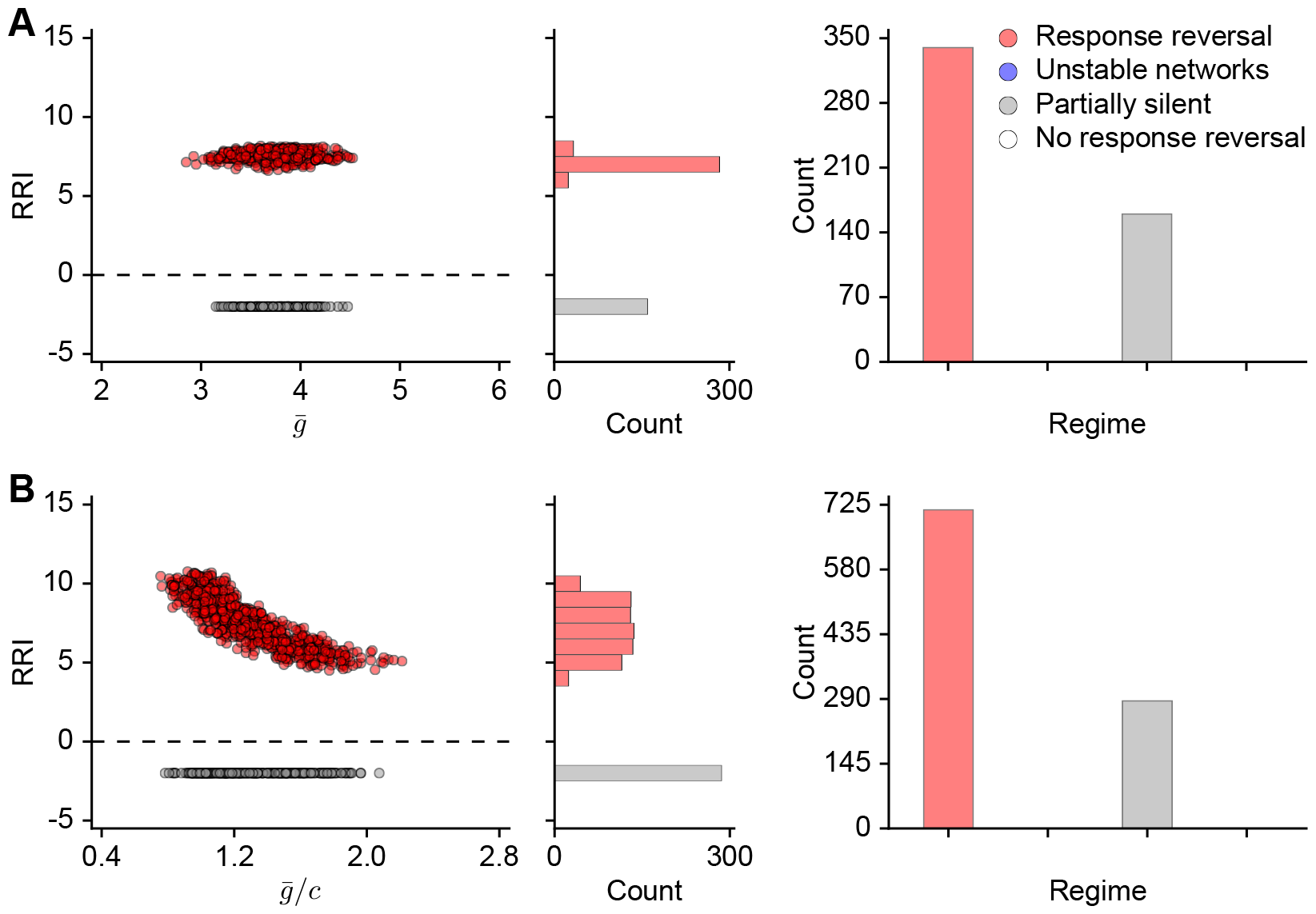
Response reversal is observed for a wide range of background and bottom-up inputs. (**A**) Left: Response Reversal Index (RRI) as a function of the mean background input 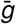. Background inputs were randomly and independently sampled from the ranges *g*_*E*_, *g*_*PV*_, *g*_*VIP*_ ∈ [3, 5], and *g*_*SST*_ ∈ [2, 4], each with a step size of 0.1. The mean background input 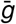 is calculated by averaging the sampled background inputs to different populations. Simulations were performed for *n* = 500 random choices. A positive RRI indicates a response reversal of SST from negative to positive, whereas a zero RRI represents no response reversal. A negative RRI indicates partially silent networks (i.e., at least one population activity is zero) or unstable networks (i.e., network activity explodes). Middle: Histogram of RRI distribution. Right: Counts of different simulation results. (**B**) Similar to A, but for the relationship between RRI and the ratio of the mean background input 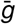 to the top-down modulatory input *c*. Additionally, *c* was randomly drawn from 2 to 4 with a step size of 0.1. Simulations were performed for *n* = 1000 random choices. All simulations were performed with top-down modulation *c* = 3.

**Fig. S12.**
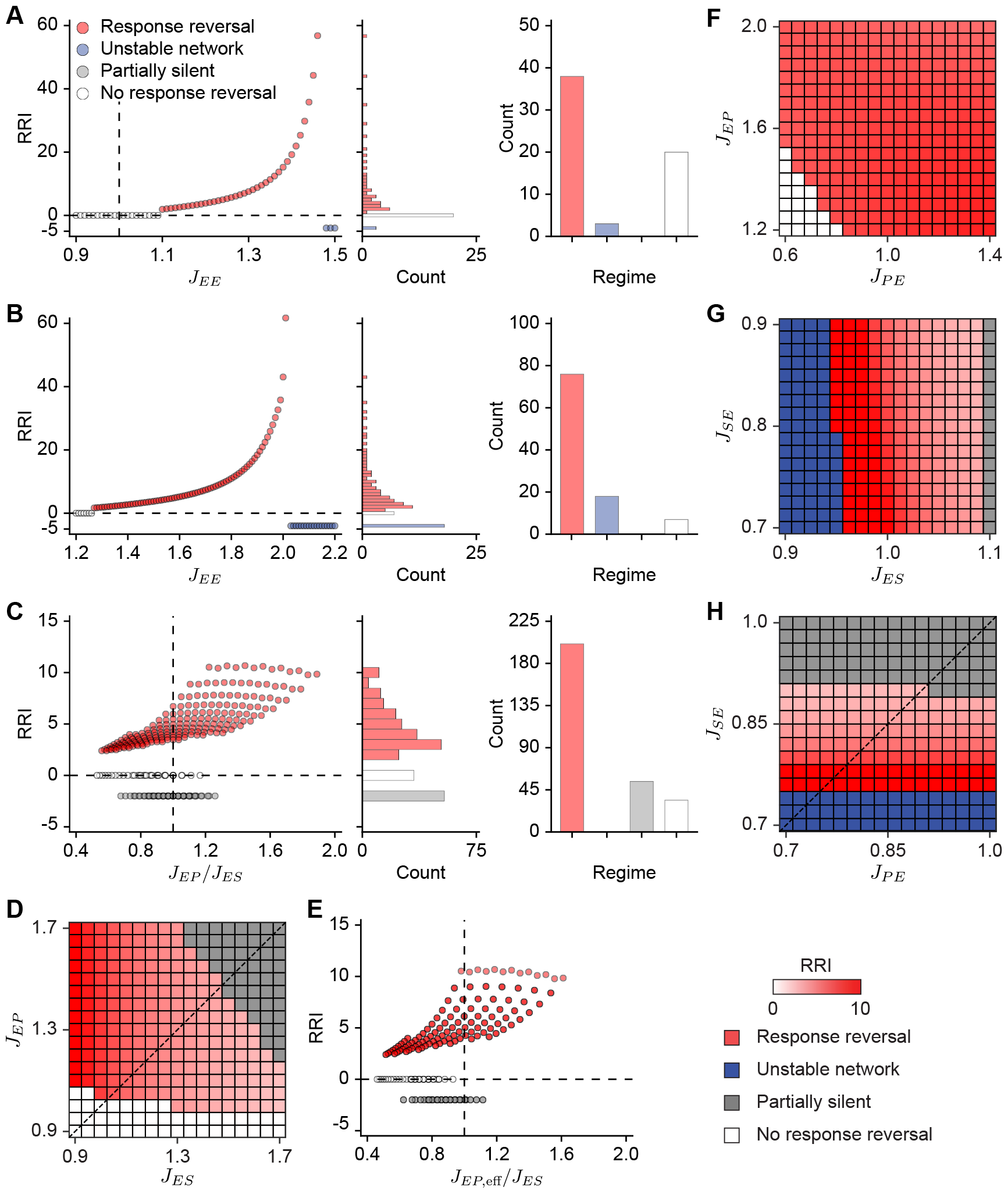
Response reversal is observed for a wide range of network connectivity. (**A**) Left: Response Reversal Index (RRI) as a function of initial weights *J*_*EE*_. A positive RRI indicates a response reversal of SST from negative to positive, whereas a zero RRI represents no response reversal. A negative RRI indicates partially silent networks (i.e., at least one population activity is zero) or unstable networks (i.e., network activity explodes). The vertical dashed line indicates *J*_*EE*_ = 1. Middle: Histogram of RRI distribution. Right: Counts of different simulation results. (**B**) Similar to A but with higher inhibitory weights (see Tab. S5). (**C**) Similar to A but for the ratio of the *J*_*EP*_ */J*_*ES*_ weights. The vertical dashed line indicates a ratio of 1. The initial values of *J*_*EP*_ and *J*_*ES*_ are taken from 0.9 to 1.7 with a stepsize of 0.05. (**D**) RRI for different initial combinations of *J*_*EP*_ and *J*_*ES*_. (**E**) Similar to C, but as a function of the effective ratio between *J*_*EP*_ and *J*_*ES*_ by multiplying the short-term depression variable *x*_*EP*_ with *J*_*EP*_. Here, *x*_*EP*_ is determined by its steady-state value before top-down modulation at the low bottom-up input (*α* = 0). (**F**) RRI for different initial weights of *J*_*PE*_ and *J*_*EP*_. (**G**) Similar to F but for different initial weights of *J*_*ES*_ and *J*_*SE*_. (**H**) Similar to F but for different initial weights of *J*_*PE*_ and *J*_*SE*_. All simulations were performed with top-down modulation *c* = 3.

**Table S1:** Parameters for networks with iSTP.

| **Network dynamics and network connectivity** |  |  |  |
| --- | --- | --- | --- |
| **Symbol** | **Value** | **Unit** | **Description** |
| $\tau_E$ | 20 | ms | time constant of E rate dynamics |
| $\tau_P$ | 10 | ms | time constant of PV rate dynamics |
| $\tau_S$ | 10 | ms | time constant of SST rate dynamics |
| $\tau_V$ | 10 | ms | time constant of VIP rate dynamics |
| $J_{EE}$ | 1.3 | a.u. | connection strength from E to E |
| $J_{EP}$ | 1.6 | a.u. | connection strength from PV to E |
| $J_{ES}$ | 1.0 | a.u. | connection strength from SST to E |
| $J_{PE}$ | 1.0 | a.u. | connection strength from E to PV |
| $J_{PP}$ | 1.3 | a.u. | connection strength from PV to PV |
| $J_{PS}$ | 0.8 | a.u. | connection strength from SST to PV |
| $J_{SE}$ | 0.8 | a.u. | connection strength from E to SST |
| $J_{SV}$ | 0.6 | a.u. | connection strength from VIP to SST |
| $J_{VE}$ | 1.1 | a.u. | connection strength from E to VIP |
| $J_{VP}$ | 0.4 | a.u. | connection strength from PV to VIP |
| $J_{VS}$ | 0.4 | a.u. | connection strength from SST to VIP |
| **Short-term plasticity** |  |  |  |
| $\tau_x$ | 100 | ms | time constant of short-term depression |
| $U_d$ | 1 | a.u. | depression factor |
| $\tau_u$ | 400 | ms | time constant of short-term facilitation |
| $U_f$ | 1 | a.u. | facilitation factor |
| $U_{\max}$ | 3 | a.u. | maximum value of the facilitation variable |
| **Inputs** |  |  |  |
| $g_E$ | 4 | a.u. | background input to E |
| $g_P$ | 4 | a.u. | background input to PV |
| $g_S$ | 3 | a.u. | background input to SST |
| $g_V$ | 4 | a.u. | background input to VIP |
| $c$ | 3 | a.u. | top-down input to VIP |

**Table S2:** Parameters for networks also including E-to-E STD.

| **Network dynamics and network connectivity** |  |  |  |
| --- | --- | --- | --- |
| **Symbol** | **Value** | **Unit** | **Description** |
| $\tau_E$ | 20 | ms | time constant of E rate dynamics |
| $\tau_P$ | 10 | ms | time constant of PV rate dynamics |
| $\tau_S$ | 10 | ms | time constant of SST rate dynamics |
| $\tau_V$ | 10 | ms | time constant of VIP rate dynamics |
| $J_{EE}$ | 1.8 | a.u. | connection strength from E to E |
| $J_{EP}$ | 2.0 | a.u. | connection strength from PV to E |
| $J_{ES}$ | 1.0 | a.u. | connection strength from SST to E |
| $J_{PE}$ | 1.4 | a.u. | connection strength from E to PV |
| $J_{PP}$ | 1.3 | a.u. | connection strength from PV to PV |
| $J_{PS}$ | 0.8 | a.u. | connection strength from SST to PV |
| $J_{SE}$ | 0.9 | a.u. | connection strength from E to SST |
| $J_{SV}$ | 0.6 | a.u. | connection strength from VIP to SST |
| $J_{VE}$ | 1.1 | a.u. | connection strength from E to VIP |
| $J_{VP}$ | 0.4 | a.u. | connection strength from PV to VIP |
| $J_{VS}$ | 0.4 | a.u. | connection strength from SST to VIP |
| **Short-term plasticity** |  |  |  |
| $\tau_x$ | 100 | ms | time constant of short-term depression |
| $\tau_x^{EE}$ | 10 | ms | time constant of E-to-E short-term depression |
| $U_d$ | 1 | a.u. | depression factor |
| $U_d^{EE}$ | 0.3 | a.u. | E-to-E depression factor |
| $\tau_u$ | 400 | ms | time constant of short-term facilitation |
| $U_f$ | 1 | a.u. | facilitation factor |
| $U_{\max}$ | 3 | a.u. | maximum value of the facilitation variable |
| **Inputs** |  |  |  |
| $g_E$ | 7 | a.u. | background input to E |
| $g_P$ | 7 | a.u. | background input to PV |
| $g_S$ | 5 | a.u. | background input to SST |
| $g_V$ | 7 | a.u. | background input to VIP |
| $c$ | 3 | a.u. | top-down input to VIP |

**Table S3:** Parameters for networks also including E-to-SST STF.

| **Network dynamics and network connectivity** |  |  |  |
| --- | --- | --- | --- |
| **Symbol** | **Value** | **Unit** | **Description** |
| $\tau_E$ | 20 | ms | time constant of E rate dynamics |
| $\tau_P$ | 10 | ms | time constant of PV rate dynamics |
| $\tau_S$ | 10 | ms | time constant of SST rate dynamics |
| $\tau_V$ | 10 | ms | time constant of VIP rate dynamics |
| $J_{EE}$ | 1.3 | a.u. | connection strength from E to E |
| $J_{EP}$ | 1.5 | a.u. | connection strength from PV to E |
| $J_{ES}$ | 0.9 | a.u. | connection strength from SST to E |
| $J_{PE}$ | 1.1 | a.u. | connection strength from E to PV |
| $J_{PP}$ | 1.3 | a.u. | connection strength from PV to PV |
| $J_{PS}$ | 0.8 | a.u. | connection strength from SST to PV |
| $J_{SE}$ | 0.5 | a.u. | connection strength from E to SST |
| $J_{SV}$ | 0.6 | a.u. | connection strength from VIP to SST |
| $J_{VE}$ | 1.1 | a.u. | connection strength from E to VIP |
| $J_{VP}$ | 0.3 | a.u. | connection strength from PV to VIP |
| $J_{VS}$ | 0.2 | a.u. | connection strength from SST to VIP |
| **Short-term plasticity** |  |  |  |
| $\tau_x$ | 100 | ms | time constant of short-term depression |
| $U_d$ | 1 | a.u. | depression factor |
| $\tau_u$ | 400 | ms | time constant of short-term facilitation |
| $U_f$ | 1 | a.u. | facilitation factor |
| $U_{\max}$ | 3 | a.u. | maximum value of the facilitation variable |
| $U_{\max}^{SE}$ | 2 | a.u. | maximum value of the E-to-SST facilitation variable |
| **Inputs** |  |  |  |
| $g_E$ | 4 | a.u. | background input to E |
| $g_P$ | 4 | a.u. | background input to PV |
| $g_S$ | 3 | a.u. | background input to SST |
| $g_V$ | 4 | a.u. | background input to VIP |
| $c$ | 3 | a.u. | top-down input to VIP |

**Table S4:** Parameters for networks with short-term plasticity on all existing connections.

| Network dynamics and network connectivity |  |  |  |
| --- | --- | --- | --- |
| Symbol | Value | Unit | Description |
| $\tau_E$ | 20 | ms | time constant of E rate dynamics |
| $\tau_P$ | 10 | ms | time constant of PV rate dynamics |
| $\tau_S$ | 10 | ms | time constant of SST rate dynamics |
| $\tau_V$ | 10 | ms | time constant of VIP rate dynamics |
| $J_{EE}$ | 1.7 | a.u. | connection strength from E to E |
| $J_{EP}$ | 2.1 | a.u. | connection strength from PV to E |
| $J_{ES}$ | 1.5 | a.u. | connection strength from SST to E |
| $J_{PE}$ | 1.0 | a.u. | connection strength from E to PV |
| $J_{PP}$ | 1.2 | a.u. | connection strength from PV to PV |
| $J_{PS}$ | 1.3 | a.u. | connection strength from SST to PV |
| $J_{SE}$ | 0.7 | a.u. | connection strength from E to SST |
| $J_{SV}$ | 0.4 | a.u. | connection strength from VIP to SST |
| $J_{VE}$ | 0.9 | a.u. | connection strength from E to VIP |
| $J_{VP}$ | 0.5 | a.u. | connection strength from PV to VIP |
| $J_{VS}$ | 0.4 | a.u. | connection strength from SST to VIP |
| Short-term plasticity |  |  |  |
| $\tau_x$ | 100 | ms | time constant of short-term depression |
| $\tau_x^{EE}$ | 10 | ms | time constant of E-to-E short-term depression |
| $U_d^{EE}$ | 0.19 | a.u. | E-to-E depression factor |
| $U_d^{EP}$ | 0.49 | a.u. | PV-to-E depression factor |
| $U_d^{ES}$ | 0.12 | a.u. | SST-to-E depression factor |
| $U_d^{PE}$ | 0.04 | a.u. | E-to-PV depression factor |
| $U_d^{PP}$ | 0.5 | a.u. | PV-to-PV depression factor |
| $U_d^{PS}$ | 0.11 | a.u. | SST-to-PV depression factor |
| $U_d^{VP}$ | 0.37 | a.u. | PV-to-VIP depression factor |
| $\tau_u$ | 400 | ms | time constant of short-term facilitation |
| $U_f^{SE}$ | 0.18 | a.u. | E-to-SST facilitation factor |
| $U_f^{VE}$ | 0.03 | a.u. | E-to-VIP facilitation factor |
| $U_f^{VS}$ | 0.28 | a.u. | SST-to-VIP facilitation factor |
| $U_f^{SV}$ | 0.05 | a.u. | VIP-to-SST facilitation factor |
| $U_{\max}$ | 3 | a.u. | maximum value of the facilitation variable |
| $U_{\max}^{SE}$ | 2 | a.u. | maximum value of the E-to-SST facilitation variable |
| Inputs |  |  |  |
| $g_E$ | 5 | a.u. | background input to E |
| $g_P$ | 5 | a.u. | background input to PV |
| $g_S$ | 3 | a.u. | background input to SST |
| $g_V$ | 5 | a.u. | background input to VIP |
| $c$ | 3 | a.u. | top-down input to VIP |

**Table S5:** Parameters for sensitivity analysis of network connectivity.

| **Network dynamics and network connectivity** |  |  |  |
| --- | --- | --- | --- |
| **Symbol** | **Value** | **Unit** | **Description** |
| $\tau_E$ | 20 | ms | time constant of E rate dynamics |
| $\tau_P$ | 10 | ms | time constant of PV rate dynamics |
| $\tau_S$ | 10 | ms | time constant of SST rate dynamics |
| $\tau_V$ | 10 | ms | time constant of VIP rate dynamics |
| $J_{EE}$ | [1.2, 2.2] | a.u. | connection strength from E to E |
| $J_{EP}$ | 1.7 | a.u. | connection strength from PV to E |
| $J_{ES}$ | 1.4 | a.u. | connection strength from SST to E |
| $J_{PE}$ | 2.2 | a.u. | connection strength from E to PV |
| $J_{PP}$ | 1.6 | a.u. | connection strength from PV to PV |
| $J_{PS}$ | 1.1 | a.u. | connection strength from SST to PV |
| $J_{SE}$ | 1.0 | a.u. | connection strength from E to SST |
| $J_{SV}$ | 0.6 | a.u. | connection strength from VIP to SST |
| $J_{VE}$ | 1.3 | a.u. | connection strength from E to VIP |
| $J_{VP}$ | 0.4 | a.u. | connection strength from PV to VIP |
| $J_{VS}$ | 0.4 | a.u. | connection strength from SST to VIP |
| **Short-term plasticity** |  |  |  |
| $\tau_x$ | 100 | ms | time constant of short-term depression |
| $U_d$ | 1 | a.u. | depression factor |
| $\tau_u$ | 400 | ms | time constant of short-term facilitation |
| $U_f$ | 1 | a.u. | facilitation factor |
| $U_{\max}$ | 3 | a.u. | maximum value of the facilitation variable |
| **Inputs** |  |  |  |
| $g_E$ | 4 | a.u. | background input to E |
| $g_P$ | 4 | a.u. | background input to PV |
| $g_S$ | 3 | a.u. | background input to SST |
| $g_V$ | 4 | a.u. | background input to VIP |
| $c$ | 3 | a.u. | top-down input to VIP |

## Notes

### Competing Interest Statement

The authors have declared no competing interest.

### Summary of Updates

Additional sections and analyses are added.

